# Coregulation by *arc*A and *fnr* protects *Salmonella* Typhimurium from bile stress by maintaining redox homeostasis and membrane integrity

**DOI:** 10.64898/2026.06.02.729527

**Authors:** Deepti Chandra, Madhulika Singh, Dipankar Nandi

## Abstract

Bile salts constitute a major antimicrobial barrier encountered by *Salmonella* Typhimurium during infections. Here, we identified the interplay of two global regulators, Fnr (fumarate and nitrate reductase) and ArcA (aerobic respiration control protein A), in determining adaptive responses to bile. We utilized WT, Δ*fnr,* Δ*arc*A and Δ*arc*AΔ*fnr* strains and compared the expression and functional responses of *S*. Typhimurium to bile. The highlights of this study are: First, the Δ*arc*A and Δ*arc*AΔ*fnr* strain form smaller colonies on LB agar plates and the Δ*arc*AΔ*fnr* strain displays lower motility. Second, qRT-PCR expression analysis demonstrates that *fnr* is induced early with bile, followed by *arc*A. Also, *arc*A transcripts are lower in the Δ*fnr* strain and *fnr* expression is also partially lower in the Δ*arc*A strain. Third, the Δ*arc*A and Δ*fnr* strains display partial sensitivity to bile, whereas the Δ*arc*AΔ*fnr* strain exhibits hypersensitivity to bile. Upon bile exposure, the Δ*arc*AΔ*fnr* strain displays elevated transcripts of the major antioxidant genes (*sod*A and *kat*G) and outer membrane protein (*omp*C), higher induction of reactive oxygen species (ROS), and greater membrane damage. Fourth, intracellular nitrite is induced earlier than ROS with bile. Fifth, *arc*A and *fnr* protects *S*. Typhimurium from bile induced stress by activating the nitrate metabolism pathway, which lowers ROS. Functionally, pretreatment of the deletion strains with sodium nitrate reduces ROS, improves membrane integrity and survival with bile. Overall, these findings demonstrate that *arc*A and *fnr* function cooperatively to utilize alternate electron acceptors, reduce dependence on aerobic respiration and lower ROS to improve survival of *S*. Typhimurium during bile stress.

## Introduction

*Salmonella enterica* belongs to the family Enterobacteriaceae and is a foodborne pathogen. *S*. Typhi causes typhoid fever, whereas non-typhoidal *Salmonella* (NTS) serovars are frequently zoonotic and can be divided into non-invasive causing gastrointestinal self-limiting infections and invasive (iNTS) leading to sepsis and meningitidis (Kumar et al., 2025; Lamichhane et al., 2024). Between 200 million and 1 billion cases of *Salmonella* infections are reported globally each year with non-typhoidal *Salmonella* causing 93 million cases of gastroenteritis and 155,000 fatalities (Kumar et al., 2025; Lamichhane et al., 2024). Salmonellosis is a major global public health concern due to its increasing prevalence as a foodborne infection, the emergence of antimicrobial resistant strains, and high mortality associated with NTS infections (Kumar et al., 2025; Lamichhane et al., 2024). NTS infections often occur in susceptible populations, including HIV infected individuals and young children with comorbidities such as malaria, anaemia, or malnutrition (Kumar et al., 2025; Tack et al., 2020). In addition, *S*. Typhi infections are corelated to gallbladder and colon cancers (Stévenin & Neefjes, 2025).

Historically, bile has been associated with *Salmonella* infections: the first identified asymptomatic carrier was Mary Mallon, a cook who spread typhoid. *Salmonella* is more resistant to bile and is often present in gall stones of typhoid carriers (Crawford et al., 2010). Bile is a potent detergent that aids in the emulsification of bile lipids and dietary lipids and bile exposure significantly affects *Salmonella* in many different ways: changes in outer membrane permeability (Ray et al., 2019), an increase in ROS (Singh et al., 2023), an increase in DNA damage (Prieto et al., 2004), and suppresses intracellular invasion (Eade et al., 2016; Giraud et al., 2024). Our laboratory has previously identified cold shock protein E (CspE) to be important in resisting bile stress (Ray et al., 2019). Subsequently, a RNA-seq analysis has shown that *S*. Typhimurium undergoes metabolic adaptation by activating genes involved in nitrate metabolism which boosts the survival of bacteria upon bile stress. This study identified the fumarate and nitrate metabolism regulator (*fnr*) to play a partial role in imparting bile tolerance (Singh et al., 2024). Fnr, an oxygen sensing transcription factor, is important in nitrate respiration and regulates the expression of genes involved in multiple pathways (Fink et al., 2007). In *S*. Typhimurium, another key sensor of the redox changes is the ArcB/ArcA two-component system, and together with Fnr, it transcriptionally regulates metabolic transition from aerobic to anaerobic metabolism. ArcB senses the redox state of the membrane quinone pool, whereas the response regulator ArcA is responsible for metabolic shift from aerobic to anaerobic conditions, controls genes related to flagella biosynthesis, cellular metabolism and motility (Evans et al., 2011; Brown et al., 2022). The Fnr and Arc system regulates metabolic adaptation during oxidative stress (Fink et al., 2007; Pardo-Esté et al., 2018). We have shown previously that bile stress results in a secondary oxidative stress within the *S*. Typhimurium and Fnr contributes to resistance (Singh et al., 2024). However, the roles of ArcA and its interaction with Fnr during bile stress remain unknown.

In this study, we examine the roles of *arc*A and *fnr* in *S*. Typhimurium during bile stress. We studied Δ*arc*A and Δ*fnr*, as well as the Δ*arc*AΔ*fnr* strains for cell growth, cell morphology, colony formation, and bile sensitivity in comparison to the WT strain. We demonstrate that *S*. Typhimurium utilizes nitrate as an alternative electron acceptor that provides growth advantage upon bile stress. To the best of our knowledge, this is the first study to characterize effects on *S*. Typhimurium with deletions in both *arcA* and *fnr*, their roles in nitrate metabolism and the interrelationship between nitrate and oxidative stress during bile stress responses. This study provides new insights on the combined roles of ArcA and Fnr regulators in the adaptation of *S.* Typhimurium to bile.

## Results

### The two-component signalling system response regulator *arc*A modulates *S.* Typhimurium bile stress response

A previous global transcriptome profile from our lab, comparing wild type (WT) *S.* Typhimurium with and without bile exposure reported that *fnr* plays a partial role in the survival of *S.* Typhimurium during bile stress (Singh et al., 2024). The partial growth defect of Δ*fnr* indicated important roles of other regulators for bacterial survival during bile treatment. The global transcriptomic compendium of *S*. Typhimurium under different stress conditions (Kröger et al., 2013) revealed higher expression of anaerobic metabolism regulators, *arc*A and *fnr*, upon bile shock (Table S6). Both *fnr* and *arc*A were induced during bile stress (Fig. S2B). This observation was supported by the previous RNA-seq from our laboratory (GEO accession number GSE248397) (Singh et al., 2024), which confirmed differential expression of genes involved in anaerobiosis (Fig. S2A). To confirm this observation, we performed a qRT-PCR analysis to examine the expression of *arc*A and *fnr* in the WT strain post bile treatment. At 3 hours, there was a significant induction in transcript levels of *fnr*; however, *arc*A transcripts were induced later, i.e. at 5 h (Fig. 1A). This indicated that *fnr* showed an early response to bile stress, whereas *arc*A displayed a delayed adaptive response.

**Fig. 1.**
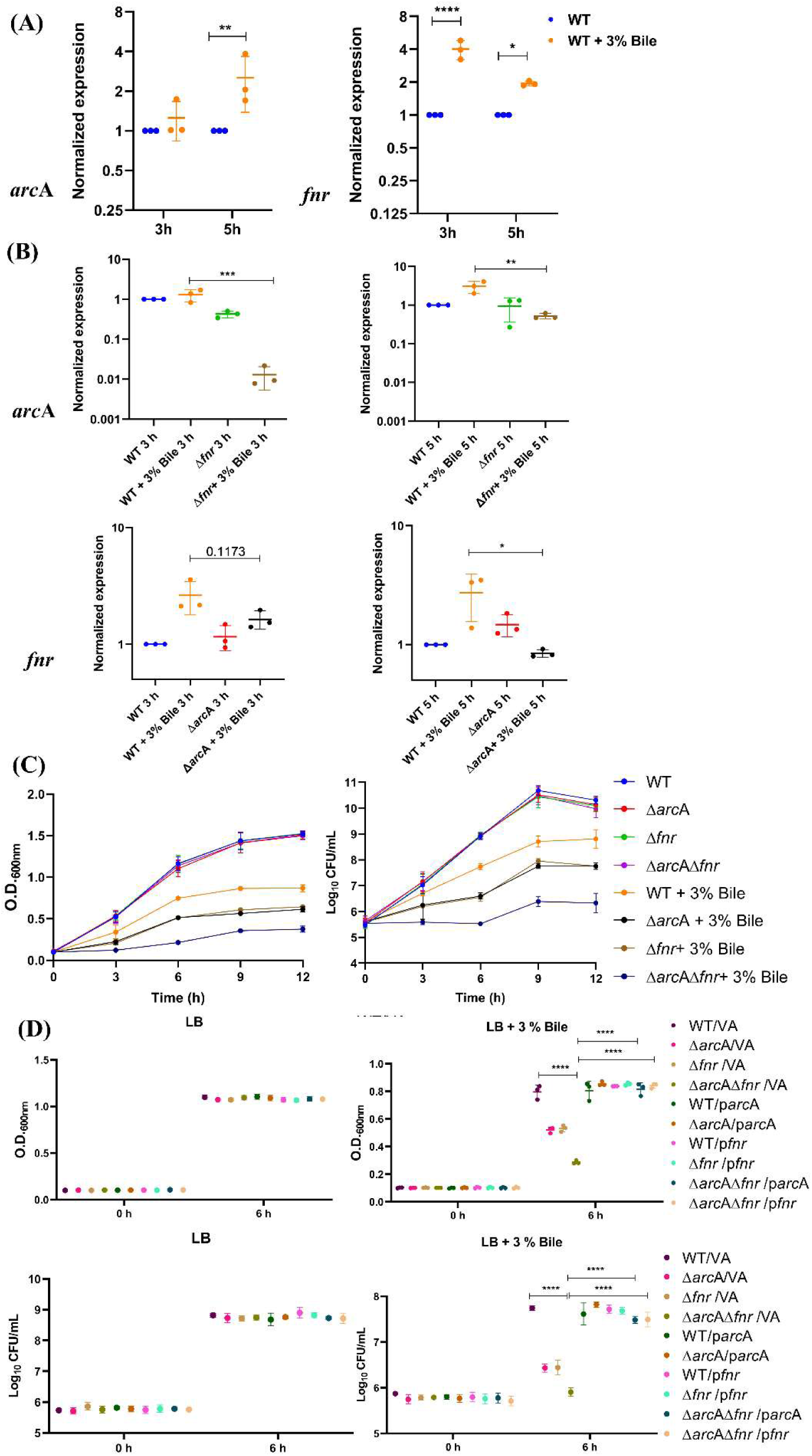
The Δ*arc*AΔ*fnr* strain shows increased sensitivity to bile stress as compared to WT and single mutants Δ*arc*A and Δ*fnr*. (A) qRT-PCR analysis of *arc*A and *fnr* in bile treated WT upon 3 h and 5 h of bile treatment. For qRT-PCR *P* values were measured by ordinary two-way ANOVA. (B) qRT-PCR analysis of *arc*A in Δ*fnr* and *fnr* in Δ*arc*A strain upon bile treatment. For qRT-PCR *P* values were measured by ordinary one-way ANOVA. The data is presented as mean ± SD from 3 independent biological experiments. \**P*< 0.05, \*\**P*< 0.01, \*\*\**P*<0.001, \*\*\*\**P* < 0.0001. (C) Growth of WT, Δ*arc*A, Δ*fnr* and Δ*arc*AΔ*fnr* strain in LB with and without the presence of 3% bile in a time dependent manner. (D) Complementation of Δ*arc*A with p*arc*A, Δ*fnr* with p*fnr* and Δ*arc*AΔ*fnr* with p*arc*A and p*fnr* upon 3% bile treatment. CFU were counted after plating cells with appropriate dilutions. The data is presented as mean ± SD from 3 independent biological experiments. *P* values were measured by two-way ANOVA. \*\*\*\**P* < 0.0001.

Despite the central roles of *arc*A and *fnr* in respiration, their contributions to bile stress adaptation are unclear. Given that both *arc*A and *fnr* showed differential expression upon bile exposure (Fig S2B), we investigated whether they function cooperatively in bile resistance. To evaluate the functional significance of *arc*A we generated an *arc*A deletion strain (Δ*arc*A) (Fig. S1). Growth curve analysis indicated that the Δ*arc*A strain showed a partial decrease in survival under bile stress compared to the WT strain (Fig. S2D). The complementation of the Δ*arc*A strain with p*arc*A rescued bile sensitivity (Fig. S2E). Previously, we had shown that the Δ*csp*E strain is highly sensitive to bile stress (Ray et al., 2019). We observed a significant increase in survival of Δ*csp*E strain upon *arc*A overexpression under bile treatment (Fig. S2C). Collectively, these findings indicate that bile exposure influences the differential expression of *arc*A in *S.* Typhimurium, and that *arc*A enhances bacterial survival during bile stress.

To elucidate the interplay between *arc*A and *fnr* under bile stress, we quantified the transcript levels of *arc*A in the Δ*fnr* strain and the transcript levels of *fnr* in the Δ*arc*A strain using qRT-PCR, with and without 3% bile exposure (Fig. 1B). It was observed that in the Δ*arc*A strain upon bile exposure, *fnr* transcript levels were similar to WT strain at 3 h; however, they were decreased partially at 5 h. On the other hand, *arc*A mRNA levels were significantly reduced in the Δ*fnr* strain upon bile treatment at 3 h. At 5 h, it increased comparatively in the Δ*fnr* strain but was still significantly less compared to the WT. These results indicate that *fnr* positively affects *arc*A expression under bile stress during early time period. Most likely, *arc*A and *fnr* regulate the expression of each other in parallel or through an indirect regulatory network upon bile stress.

### The Δ*arc*AΔ*fnr* strain displays increased sensitivity to bile treatment

The expression results motivated us to create a double deletion strain (Δ*arc*AΔ*fnr*) (Fig. S1). A growth analysis in the presence of different concentrations of bile (1%, 3%, 5%) demonstrated that, single mutants (Δ*arc*A and Δ*fnr*) were partially sensitive, whereas Δ*arc*AΔ*fnr* was highly sensitive to all concentrations of bile during aerobic growth (Fig. S2F). Growth kinetics of WT, Δ*arc*A, Δ*fnr* and Δ*arc*AΔ*fnr* strains showed that the Δ*arc*AΔ*fnr* cells exhibited decreased growth as compared to WT and single deletion strains (Δ*arc*A and Δ*fnr*) upon treatment with 3% bile in aerobic growth, as reflected by both O.D. and CFU (Fig. 1C). Complementation with p*arc*A and p*fnr* rescued the bile sensitive phenotype of Δ*arc*A and Δ*fnr* strains respectively. Also, the complementation with p*arc*A or p*fnr* rescued the bile hypersensitive phenotype of the Δ*arc*AΔ*fnr* strain (Fig. 1D). Stress specificity was evaluated by studying the susceptibility of Δ*arc*A, Δ*fnr*, and Δ*arc*AΔ*fnr* strains to common environmental stresses such as osmotic and temperature stress, which are also known to induce oxidative stress (Gunasekera et al., 2008; Kim et al., 2020; Marcén et al., 2017). No differences were observed between WT, Δ*arc*A, Δ*fnr*, and Δ*arc*AΔ*fnr* strains under osmotic stress conditions (Fig. S3A) or elevated temperature (42°C), and reduced temperature (25°C) compared to growth at 37°C. The Δ*arc*A and Δ*fnr* strains showed partial sensitivity and Δ*arc*AΔ*fnr* strain showed hypersensitivity to bile at both 25°C and 42°C (Fig. S3B). These results demonstrate that the responses of *fnr* and *arc*A are specific to some stress conditions, e.g. bile.

Under microaerobic growth conditions, the Δ*arc*A, Δ*fnr* strains and the Δ*arc*AΔ*fnr* double deletion strain showed reduced growth as compared to WT cells in the absence of bile (Fig. S4A). This confirmed the microaerobic sensitivity of Δ*arc*A, Δ*fnr* strains and that the sensitivity was exacerbated in the Δ*arc*AΔ*fnr* strain. Upon adjustment of the initial inoculum volumes under microaerobic growth conditions, a comparable growth of all the strains was observed in LB media. Upon bile treatment, Δ*arc*A and Δ*fnr* displayed partial sensitivity whereas Δ*arc*AΔ*fnr* was most susceptible to bile-mediated killing, exhibiting a trend similar to that observed under aerobic condition (Fig. S4B). Further, complementation with p*arc*A and p*fnr* rescued the bile sensitive phenotype of Δ*arc*A, Δ*fnr* and Δ*arc*AΔ*fnr* cells (Fig. S4C and Fig. S4D). As the results obtained were showing similar trends of bile sensitivity in aerobic and microaerobic growth conditions, we have limited this study to aerobic growth upon bile treatment.

### Δ*arc*A and Δ*arc*AΔ*fnr* displays smaller colony size on LB agar plate

We observed no significant differences in growth between the WT, Δ*arc*A, Δ*fnr* and Δ*arc*AΔ*fnr* strains when cultured in LB broth (Fig. 1C). However, when grown on LB agar plates, striking differences in colony morphology were observed. WT and Δ*fnr* colonies were comparable, whereas the Δ*arc*A strain formed smaller colonies. The Δ*arc*AΔ*fnr* also displayed a reduction in colony size similar to the Δ*arc*A strain (Fig. 2A–B). Complementation with p*arc*A restored colony size in both Δ*arc*A and Δ*arc*AΔ*fnr* strains (Fig. S5A–B). Next, we studied the bacterial cell size using atomic force microscopy on cells cultured in LB broth and on LB agar plates. Cells grown on LB agar plates showed reduced cell length relative to cells grown in LB broth; however, no strain specific differences were observed (Fig. 2C-D). These findings indicate that *arc*A impacts the colony size of *S*. Typhimurium on LB agar media.

**Fig. 2.**
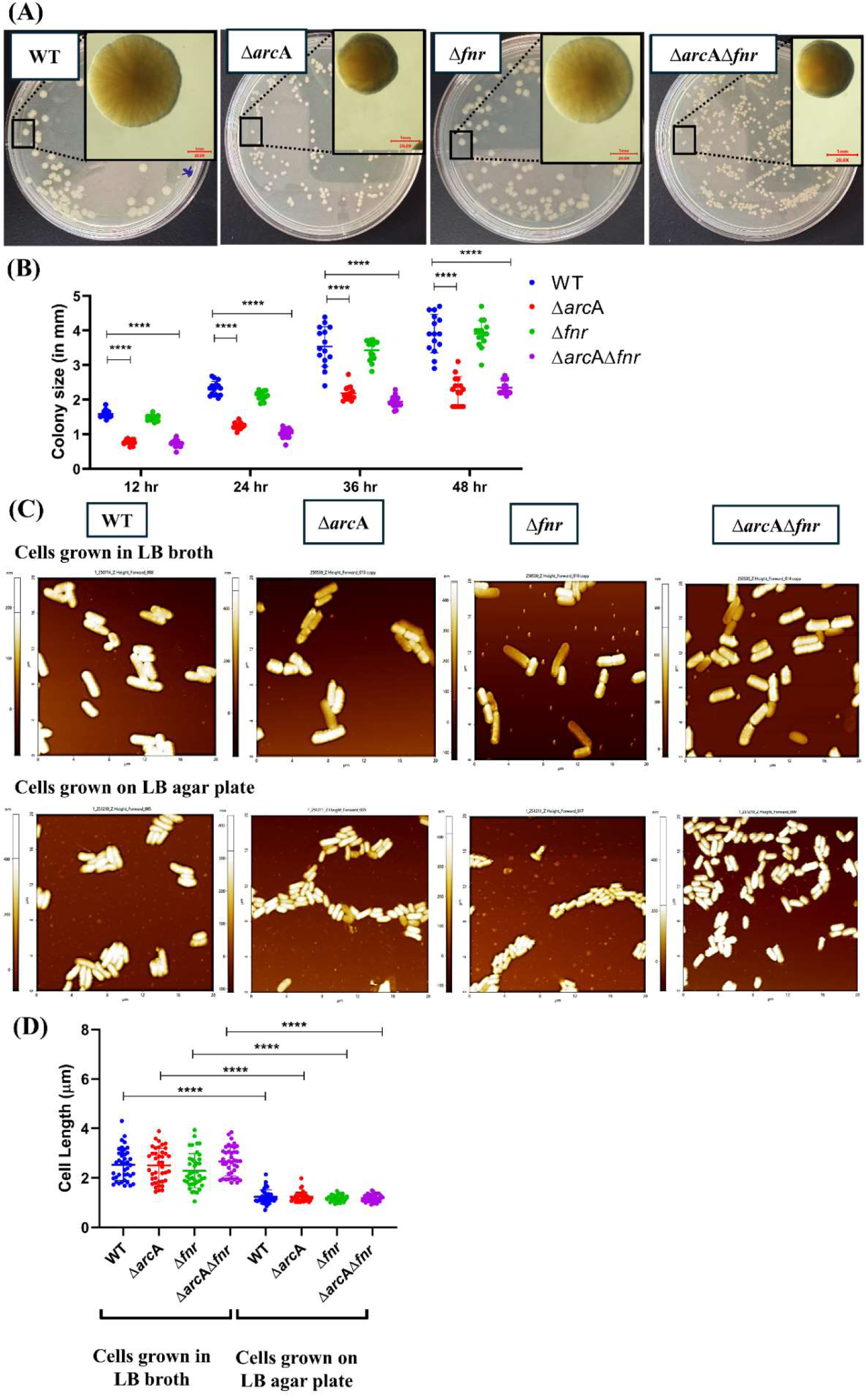
The Δ*arc*A and Δ*arc*AΔ*fnr* strains demonstrates reduced colony size. (A) Representative images of WT, Δ*arc*A, Δ*fnr* and Δ*arc*AΔ*fnr* individual colonies morphology on LB plates at 20x magnification at 36 h from 3 independent plates per strain. (B) Quantitation of the diameter of colonies. (C) The bacterial strains were grown for 6 h in LB broth (top panel) and for 12 h on LB agar plate (bottom panel), and AFM images were acquired in the non-contact mode. (D) Quantitation of cellular length of each strain was performed. For the colony size quantitation data is representative of 40 cells analysed from 3 individual experiments and presented as mean ± SD. For the cell length quantitation, the data is representative of 3 different experiments and presented as mean ± SD. *P* values were measured by two-way ANOVA using Turkey’s multiple comparison test. \*\*\*\**P* < 0.0001.

### *arc*A and *fnr* regulate the swimming and swarming motility in *S*. Typhimurium

As we had observed differences in colony size in Δ*arc*A and Δ*arc*AΔ*fnr*, we next investigated whether loss of Δ*arc*A and Δ*fnr* also affected bacterial motility. We studied the transcript levels of class I (*flh*D), class II (*fli*A, *fli*L) and class III (*mot*A) flagellar motility genes by qRT-PCR. Transcript levels of *flhD* were significantly reduced in Δ*arc*A, whereas Δ*fnr* displayed lower transcripts of *flhD fliA* and *fliL* as compared to WT. On the other hand, the Δ*arc*AΔ*fnr* strain showed lower transcript of *flh*D, *fli*A, *fli*L and *mot*A as compared to WT strain (Fig. 3A). We found that the swimming motility of Δ*arc*AΔ*fnr* strain exhibited higher decrease than the single mutants, as evidenced by markedly smaller swimming diameters compared to WT, Δ*arc*A and Δ*fnr* strains at both 8 and 16 h (Fig. 3B,D). Swarming motility was also impaired in Δ*arc*A, Δ*fnr* and Δ*arc*AΔ*fnr* strains, with the strongest reduction in swarming observed in Δ*arc*AΔ*fnr* strain at both 8 h and 16 h (Fig. 3C,D). Collectively, these results establish that *arc*A and *fnr* play a critical role in regulating motility in *S.* Typhimurium. Although motility is significantly reduced in the Δ*arc*AΔ*fnr* strain, residual movement is still observed, indicating impaired rather than abolished flagellar function.

**Fig. 3.**
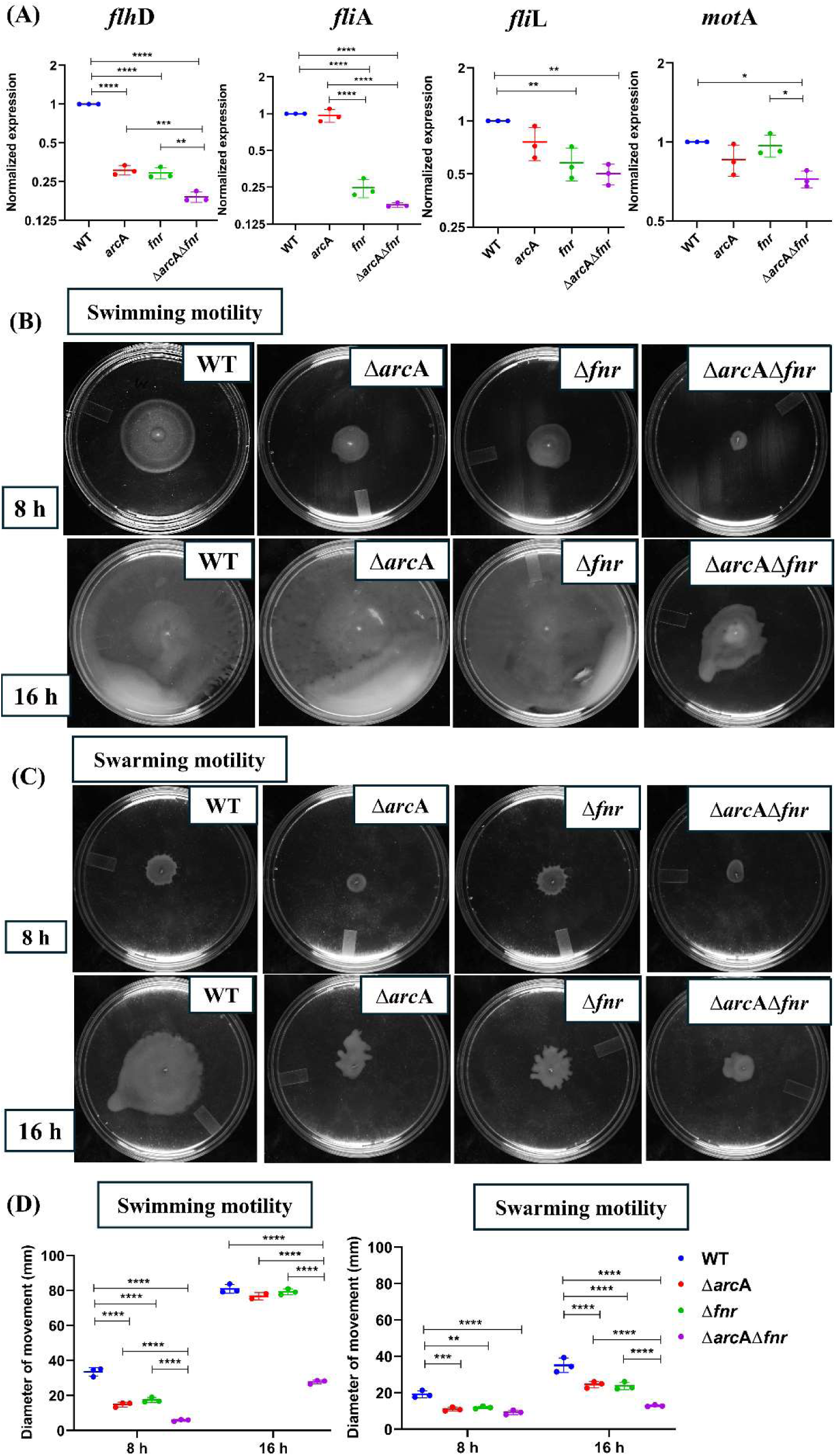
The Δ*arc*AΔ*fnr* strain exhibits severe motility defects compared to WT and single mutant Δ*arc*A and Δ*fnr* strains. (A) Transcript levels of class I (*flh*D), II (*fli*A, *fli*L) and III (*mot*A) of the flagellar regulons were analysed by qRT-PCR analysis. Cells from 6.5 h of swimming plate of all strains were used. In all panels values of WT are normalized to one and fold difference was compared. Images are representative of ≥3 independent plates per strain and time point and presented as mean ± SD. *P* values were measured by ordinary one-way ANOVA using Turkey’s multiple comparison test. \**P* <0.05, \*\**P*< 0.01, \*\*\**P*<0.001, \*\*\*\**P* < 0.0001. (B) Swimming (0.3 % w/v agar), and (C) Swarming (0.5% w/v agar) motility of indicated strains at 8 h and 16 h are shown. (D) Quantitation of diameter of the swimming and swarming motility for the indicated strains were obtained. The data is representative of 3 individual experiments and presented as mean ± SD. *P* values were measured by two-way ANOVA using Turkey’s multiple comparison test. \*\**P*< 0.01, \*\*\**P*<0.001, \*\*\*\**P* < 0.0001.

### Δ*arc*AΔ*fnr* shows increase in levels of outer membrane proteins (OMP) and membrane damage upon bile treatment

Bile stress is known to affect the expression of genes involved in the envelope stress response (Ray et al., 2019). Therefore, we studied the transcript levels of major outer membrane protein C in the four strains upon bile exposure by qRT-PCR. Upon bile stress the transcript levels of *omp*C were induced in Δ*arc*A and Δ*arc*AΔ*fnr* strain (Fig. 4A). This indicated a connection between the bile-sensitivity of the Δ*arc*A, Δ*fnr* and Δ*arc*AΔ*fnr* strains with porin mRNA amounts during bile stress. An increased amount of *omp*C might result in an increased entry of bile due to cell being more porous. This was functionally tested using the hydrophobic fluorescent probe, NPN (1-N-phenyl-1-naphthylamine) which is frequently used to evaluate the permeabilization of outer membrane of Gram-negative bacteria (Mikheyeva et al., 2023; Richard et al., 2023). The fluorescence of NPN dye increased in Δ*arc*A and Δ*fnr* strains upon bile exposure with most elevated fluorescence observed in the Δ*arc*AΔ*fnr* strain (Fig. 4B-C). The intracellular accumulation of the DNA staining dye, bisbenzimide H33258, was also studied in the four strains upon bile exposure. Correlating with the previous data, the single mutant strains Δ*arc*A and Δ*fnr* displayed an increased fluorescence of bisbenzimide H33258 with the highest fluorescence observed in the Δ*arc*AΔ*fnr* strain, upon bile exposure (Fig. S6A,B). Complementation with p*arc*A and p*fnr* in Δ*arc*A, Δ*fnr* and Δ*arc*AΔ*fnr* strains rescued the elevated fluorescence of bisbenzimide H33258 (Fig. S6C). These results indicate that there is an increase in membrane permeabilization in the Δ*arc*AΔ*fnr* strain upon bile exposure.

**Fig. 4.**
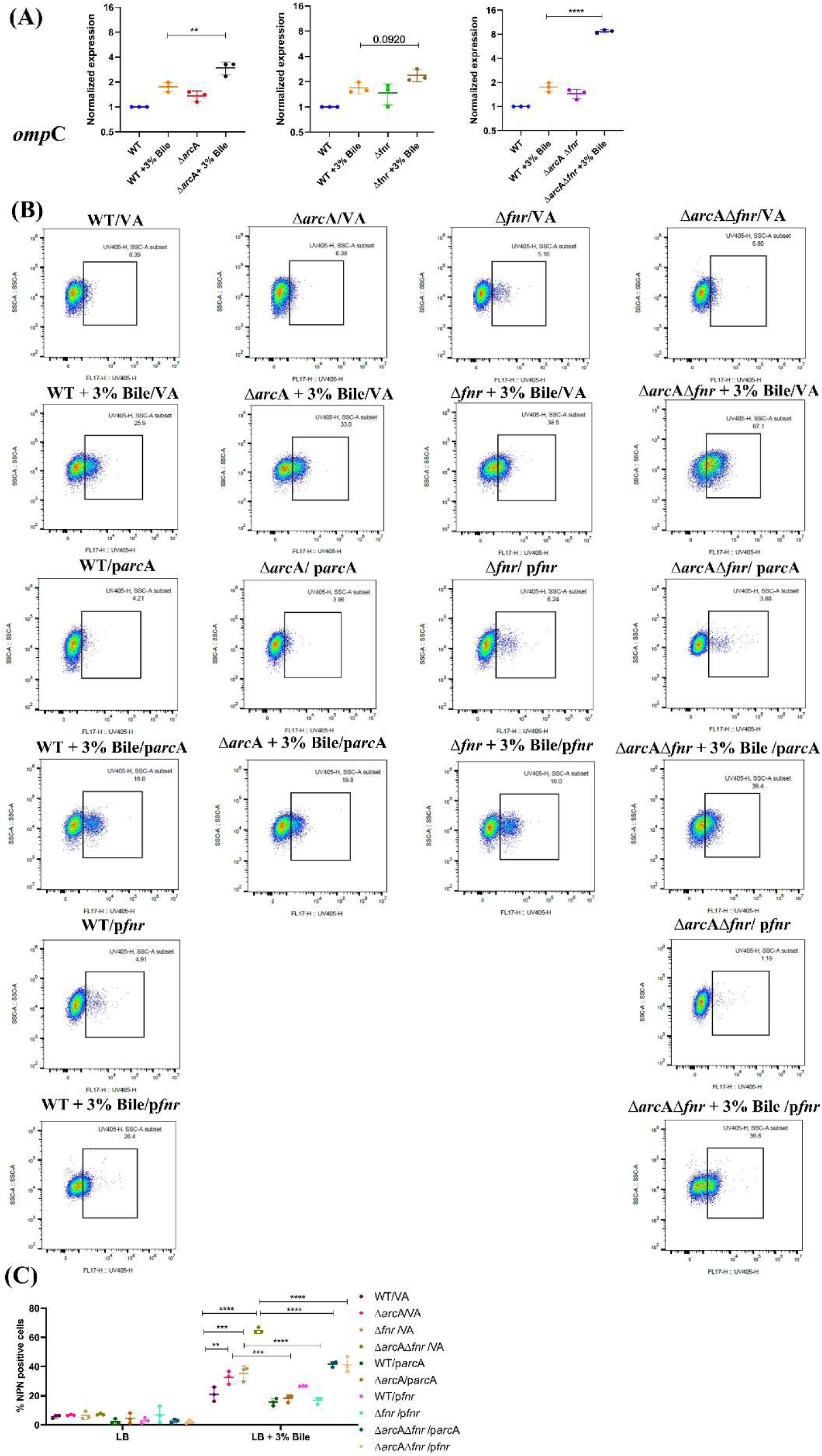
Bile salt treated Δ*arc*AΔ*fnr* strain displays increased membrane damage compared to WT and single mutant Δ*arc*A and Δ*fnr* strains. (A) Transcript levels of *omp*C were analysed by qRT-PCR analysis. Cells were harvested after 3.5 h of bile exposure. Values of WT are normalized to one and fold difference was compared. The data is representative of 3 individual experiments and presented as mean ± SD. *P* values were measured by ordinary one-way ANNOVA using Turkey’s multiple comparison test. \*\**P*< 0.01, \*\*\*\**P* < 0.0001. (B) Cells were stained with NPN and analysed by flow cytometry post 6 h of bile treatment. (C) The percentage NPN positive cells based on flow cytometry data. Data is representative of 3 individual experiments and presented as mean ± SD. *P* values were measured by two-way ANOVA using Turkey’s multiple comparison test. \*\**P*< 0.01, \*\*\**P*<0.001, \*\*\*\**P* < 0.0001.

### Δ*arc*AΔ*fnr* displays higher intracellular ROS amounts as compared to WT and single mutants upon bile treatment

Next, we studied the amounts of intracellular ROS using DCFDA staining as bile is known to induce ROS production in cells (Singh et al., 2023). qRT-PCR analysis revealed that the Δ*arc*A and Δ*fnr* strains displayed increased amounts of transcripts of antioxidant genes: *sodA* which converts O₂·^−^ to H₂O₂ and *kat*G which detoxifies H₂O₂ to H₂O and O₂. The highest amount of transcripts was observed in the Δ*arc*AΔ*fnr* strain upon treatment with bile as compared to the WT strain (Fig. 5A). Concomitantly, intracellular ROS concentrations were significantly higher in the Δ*arc*AΔ*fnr* strain than in WT and Δ*arc*A and Δ*fnr* single mutants as observed by DCFDA staining (Fig. 5B). Total ROS levels were compared between bile treated deletion strains complemented with p*arc*A and p*fnr*, which showed, lower ROS amounts (Fig. 5C), establishing that both *arc*A and *fnr* mitigate ROS generated with bile treatment. This observation was confirmed in a stress assay which showed that the double mutant was highly sensitive to hydrogen peroxide in a dose dependent manner, with significantly lower survival as compared to WT, Δ*arc*A and Δ*fnr* (Fig. 5D). Complementation with p*arc*A and p*fnr* rescued the H₂O₂ sensitive phenotype of Δ*arc*A, Δ*fnr* and Δ*arc*AΔ*fnr* strain (Fig. S7A-B). Together, these findings demonstrate the roles of *arc*A and *fnr* in defence against bile and oxidative stress.

**Fig. 5.**
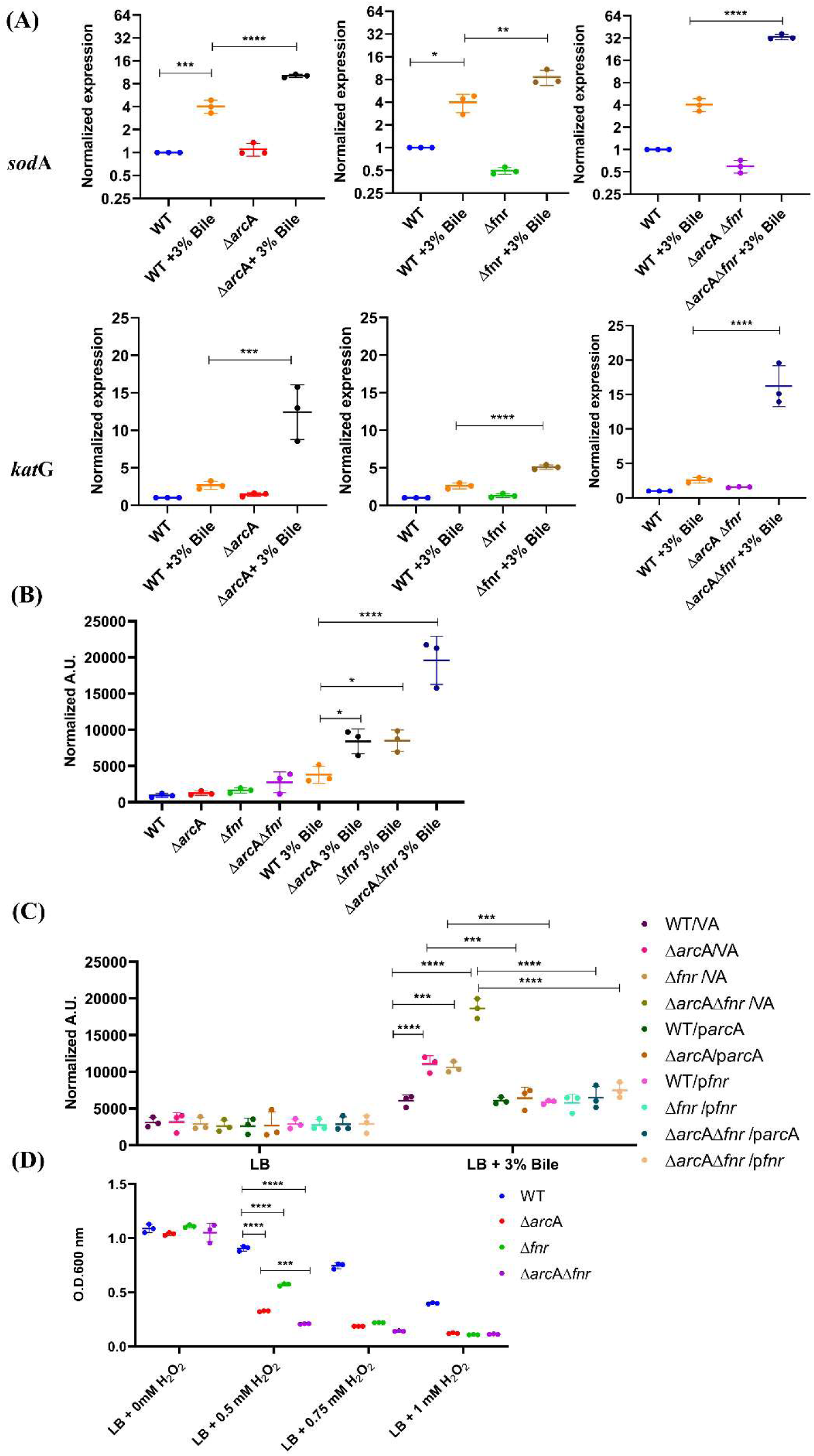
The Δ*arc*AΔ*fnr* strain exhibits elevated ROS levels and profoundly reduced survival during peroxide stress. (A) qRT-PCR analysis of antioxidant genes *sod*A and *kat*G in cells treated with bile for 3.5 h. For both genes the values of WT were normalized to one and fold difference was compared Values of WT are normalized to one and fold difference was compared. The data is representative of 3 individual experiments and presented as mean ± SD. *P* values were measured by ordinary one-way ANNOVA using Turkey’s multiple comparison test. \**P*<0.05, \*\**P*<0.01, \*\*\**P*<0.001, \*\*\*\**P*<0.0001. (B) Quantitation of intracellular ROS in WT, Δ*arc*A, Δ*fnr* and Δ*arc*AΔ*fnr* using DCFDA staining at 5 h. The data is representative of 3 individual experiments and presented as mean ± SD. *P* values were measured by ordinary one-way ANOVA using Turkey’s multiple comparison test. \**P* <0.05, \*\**P*< 0.01, \*\*\**P*<0.001, \*\*\*\**P* < 0.0001. (C) DCFDA staining was used to quantify cellular ROS, and fluorescence was normalized to OD_600_ to account for differences in cell density. (D) The O.D. was measured after H₂O₂ treatment in a dose dependent manner at 6 h. Data is representative of 3 independent biological experiments and presented as mean ± SD. *P* values were measured by two-way ANOVA using Turkey’s multiple comparison test. \*\*\**P*<0.001, \*\*\*\**P* < 0.0001.

### Nitrate pretreatment restores the growth of the Δ*arc*AΔ*fnr* strain upon bile exposure

A previous RNA-seq analysis from our laboratory had demonstrated that genes involved in nitrate metabolism are differentially expressed in *S*. Typhimurium upon bile stress (Singh et al., 2024). Based on this, we evaluated the role of different electron acceptors during bile stress by pretreating WT, Δ*arc*A, Δ*fnr*, and Δ*arc*AΔ*fnr* strains with sodium nitrate, sodium nitrite, sodium fumarate, and dimethyl sulfoxide (DMSO). Pretreatment of cells with sodium nitrate, sodium fumarate and DMSO for 1.5h before bile exposure rescued the bile sensitive phenotype of all the strains (Fig. S8A-D). Among all electron acceptors, we focused on nitrate metabolism in this study as nitrate is the most energetically favourable electron acceptor after oxygen, having the highest standard redox potential, available to *S.* Typhimurium in the inflamed gut. Therefore, it is preferentially utilized over alternate electron acceptors (Rivera-Chávez & Bäumler, 2015).

Next, we studied the transcript levels of nitrate reductases *nar*G (*nar*GHJI operon) and *nar*L (part of the two component system *nar*XL). The proteins NarX and NarL constitute a conserved two-component regulatory system. NarX is a sensor kinase that responds to nitrate and nitrite ligands to initiate autophosphorylation and activation of the NarL response-regulator receiver domain (Mangalea & Borlee, 2022). Phosphorylated NarL is the direct activator of *narGHJI* operon which encodes for the nitrate reductase A, which suggests a positive regulation of *narG* operon (nitrate reductase) by NarL (Lopez et al., 2015; Mangalea & Borlee, 2022) . The qRT-PCR analysis indicated decreased transcript level of *nar*G and *nar*L in Δ*arc*A and Δ*fnr* with the maximum decrease observed in the Δ*arc*AΔ*fnr* strain upon bile treatment (Fig. 6A). Since nitrate is reduced to nitrite as an intermediate during nitrate respiration, intracellular nitrite estimation can be used to serve as indicator of changes in nitrate metabolism in cells (Singh et al., 2024; Tiso & Schechter, 2015). The quantitation of intracellular nitrite in WT, Δ*arc*A, Δ*fnr* and Δ*arc*AΔ*fnr* strain upon bile treatment revealed that WT had a significantly higher intracellular nitrite levels as compared to Δ*arc*A, Δ*fnr* and Δ*arc*AΔ*fnr* strains (Fig. 6B). This indicates that WT activates a robust nitrate metabolism pathway, unlike the deletion strains, that converts of NO_3_^-^ to NO_2_^-^. Additionally, pretreatment of cells with sodium nitrate (NaNO_3_) significantly increased the growth of all the strains during bile stress (Fig. 6C) These results indicate that alternate electron acceptors like fumarate, DMSO and nitrate aid the survival of *S*. Typhimurium during bile stress. This could be attributed to the metabolic shift that makes use of the available alternate electron acceptors, thereby reducing dependence on aerobic metabolism and decreasing the resultant ROS. We have tested this hypothesis in next section. Importantly, we identified a previously unrecognized function of ArcA in mitigating bile induced stress via nitrate metabolism.

**Fig. 6.**
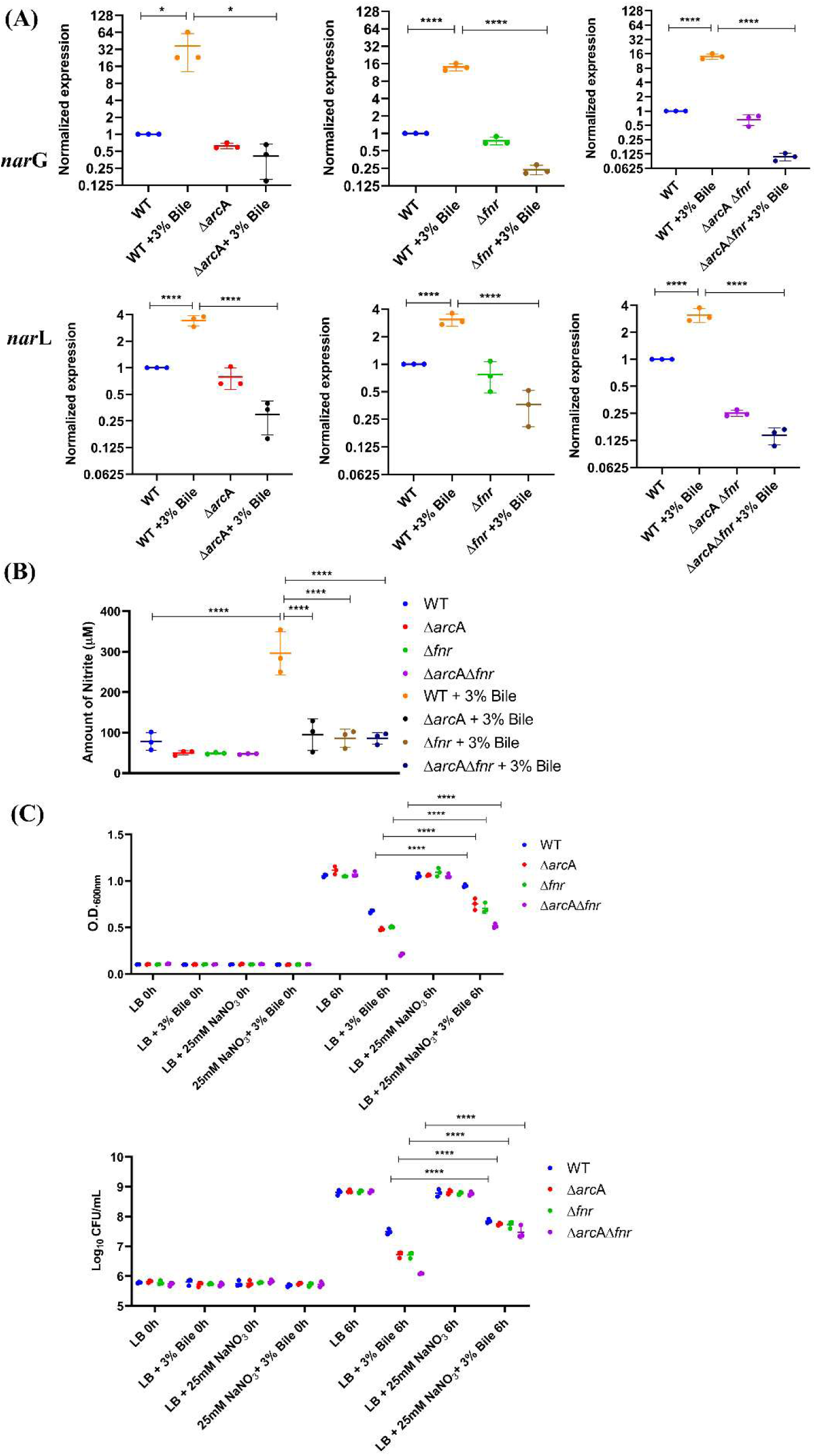
Pretreatment with nitrate enhances the survival of the Δ*arc*AΔ*fnr* stain upon bile stress. (A) Cells were harvested after 3.5 h of bile treatment and transcript levels of *nar*G and *nar*L were quantified. For both genes the values of WT were normalized to one and fold difference was compared. The data is representative of 3 individual experiments and presented as mean ± SD. *P* values were measured by ordinary one-way ANOVA using Turkey’s multiple comparison test. \*\*\*\**P* < 0.0001. (B) Quantitation of intracellular nitrite was performed at 3.5 h The data is representative of 3 individual experiments and presented as mean ± SD. *P* values were measured by ordinary one-way ANOVA using Turkey’s multiple comparison test. \*\*\*\**P* < 0.0001. (C) O.D. and CFU of cells pretreated with sodium nitrate and then exposed to 6 h of bile. The data is representative of 3 individual experiments and presented as mean ± SD. *P* values were measured by two-way ANOVA using Turkey’s multiple comparison test. \*\*\**P*<0.001, \*\*\*\**P* < 0.0001.

### Pretreatment with sodium nitrate reduces the total ROS levels and the membrane permeability upon bile stress

To better understand the mechanism by which pretreatment with sodium nitrate rescued the bile sensitivity in these strains, we studied the total ROS amounts and membrane damage. Pretreatment of cells with sodium nitrate before bile exposure decreased ROS amounts in Δ*arc*A, Δ*fnr* and Δ*arc*AΔ*fnr* strains. Although the highest degree of reduction in ROS was observed in Δ*arc*AΔ*fnr* strain, it was still higher than the WT (Fig. 7A). Also, decreased fluorescence of NPN (Fig. 7B) (Fig. S9A) and bisbenzimide H33258 (Fig. S9 B) was observed upon pretreatment with sodium nitrate before bile exposure in all strains. The Δ*arc*AΔ*fnr* strain showed a conspicuous reduction in fluorescence, indicating lower membrane damage and an almost complete recovery during bile stress. The decrease in total ROS amounts and membrane permeability explains the rescue in the bile sensitivity observed in Δ*arc*A, Δ*fnr* and Δ*arc*AΔ*fnr* strains upon exposure to sodium nitrate (Fig. 6C).

**Fig. 7.**
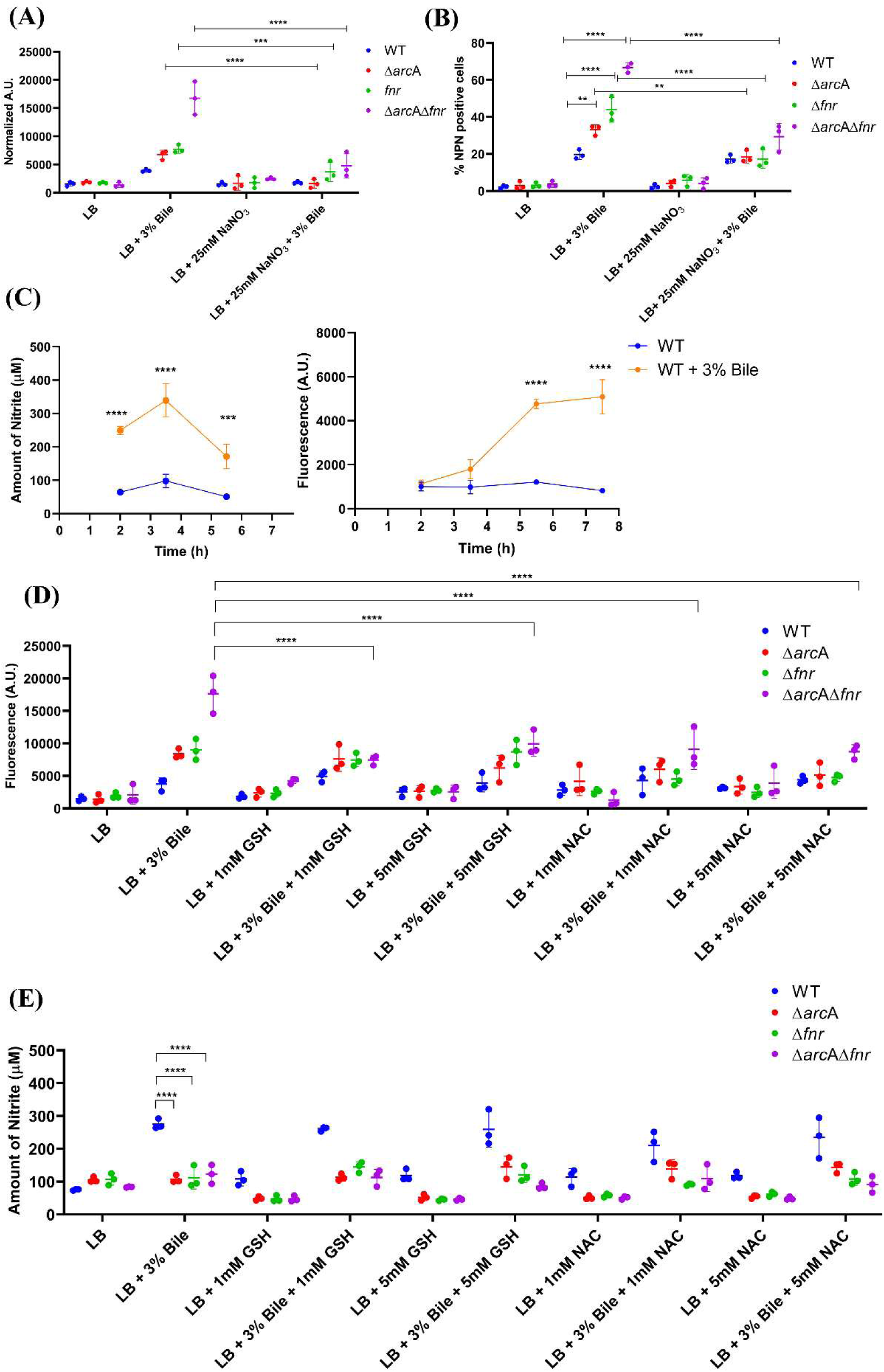
Pretreating cells with nitrate lowers ROS amounts and rescues the membrane damage in the Δ*arc*AΔ*fnr* strain treated with bile. (A) Estimation of intracellular ROS by DCFDA staining of cells treated with bile for 5 h post pretreatment with sodium nitrate. DCFDA fluorescence was normalized to OD600 and expressed as arbitrary units (A.U.). (B) The percentage NPN positive cells were plotted based on flow cytometry data. (C) Estimation of intracellular nitrite and ROS levels in WT with and without bile treatment in a time dependent manner. The \*\*\**P* < 0.001, \*\*\*\**P* < 0.0001 is the comparison between the bile treated and untreated samples at respective time points. (D) Cells were pretreated with glutathione (GSH) and N-Acetyl Cysteine (NAC) for 90 min and were treated with 3% bile for 6 h. ROS amounts were quantified by DCFDA staining after pretreating with GSH and (E) NAC. (E) Intracellular nitrite levels were quantified after pretreating with GSH and NAC. Data is representative of 3 individual experiments and presented as mean ± SD. *P* values were measured by two-way ANOVA using Turkey’s multiple comparison test. \*\**P* < 0.01, \*\*\**P* < 0.001, \*\*\*\**P* < 0.0001.

To understand the interplay between oxidative stress and nitrate metabolism upon bile exposure, we studied the ROS and nitrite amounts in the WT strain in a kinetic manner. Post-bile treatment, nitrite levels peaked early at 3.5 h; however, the ROS accumulation was maximum at 5.5 h, when the nitrite levels had dropped (Fig. 7C). This indicated that nitrate metabolism negatively impacted the total cellular ROS amounts during bile treatment. To understand this aspect better, we employed antioxidants, e.g. glutathione and N-acetyl cysteine (NAC). Pretreatment of WT, Δ*arc*A, Δ*fnr* and Δ*arc*AΔ*fnr* strains with glutathione and N-acetyl cysteine (NAC) reduced the total ROS levels in Δ*arc*AΔ*fnr* (Fig. 7D) and partially rescued the growth defect in the deletion strains during bile stress (Fig. S10A-B). Pretreatment with antioxidants did not alter the nitrite levels in WT. Interestingly, treatment with antioxidants did not increase the nitrite levels in Δ*arc*A, Δ*fnr* and Δ*arc*AΔ*fnr* strains (Fig. 7E), despite our earlier observation that pre-treatment with nitrate increased growth in Δ*arc*A, Δ*fnr* and Δ*arc*AΔ*fnr* strains (Fig. S10A-B) and lowered ROS amounts in Δ*arc*AΔ*fnr* (Fig. 7A). Overall, these results demonstrated that nitrate metabolism negatively correlates with ROS during bile stress; in contrast, ROS scavenging failed to decrease nitrite levels. We have shown that nitrite levels peak before ROS accumulation in bile-treated cells and even after a decline, is sustained at levels higher than the untreated cells (Fig. 7C). Therefore, nitrate metabolism possibly is an adaptive mechanism to mitigate the deleterious effects of bile stress-induced oxidative damage.

## Discussion

Bile exposure presents a multifactorial challenge to *S*. Typhimurium by inducing a variety of responses such as repression in cell invasion, higher DNA damage and induction of SOS responses, increased membrane damage and ROS production (Eade et al., 2016; Giraud et al., 2024; Prieto et al., 2004; Ray et al., 2019; Singh et al., 2023). Previously, our laboratory had performed RNA sequencing between WT with and without bile treatment to study the global transcriptome profile of *S*. Typhimurium and found that upon challenge with bile the bacteria undergo major metabolic remodelling (Singh et al., 2024). In this study, we used Δ*fnr* and generated Δ*arc*A and Δ*arc*AΔ*fnr* strains and studied their roles in colony morphology, motility, redox metabolism, membrane permeability and nitrate metabolism in survival upon bile stress. The Δ*arc*AΔ*fnr* strain exhibits hypersensitivity to bile treatment and complementation with p*arc*A and p*fnr* rescued the bile sensitive phenotype of this strain (Fig. 1D), indicating that *arc*A and *fnr* work together to promote bile resistance. Crucially, the lack of similar defects under temperature or osmotic stress emphasizes that *arc*A and *fnr* are specifically sensitive to bile stress rather than other stresses studied (Fig. S3). Our data reveal that *arc*A and *fnr* play key roles in maintaining cellular homeostasis during bile exposure by utilizing nitrate as an alternative electron acceptor, and that their combined loss leads to severe bile sensitivity due to increased ROS and increased membrane permeability (Fig. 8).

**Fig. 8.**
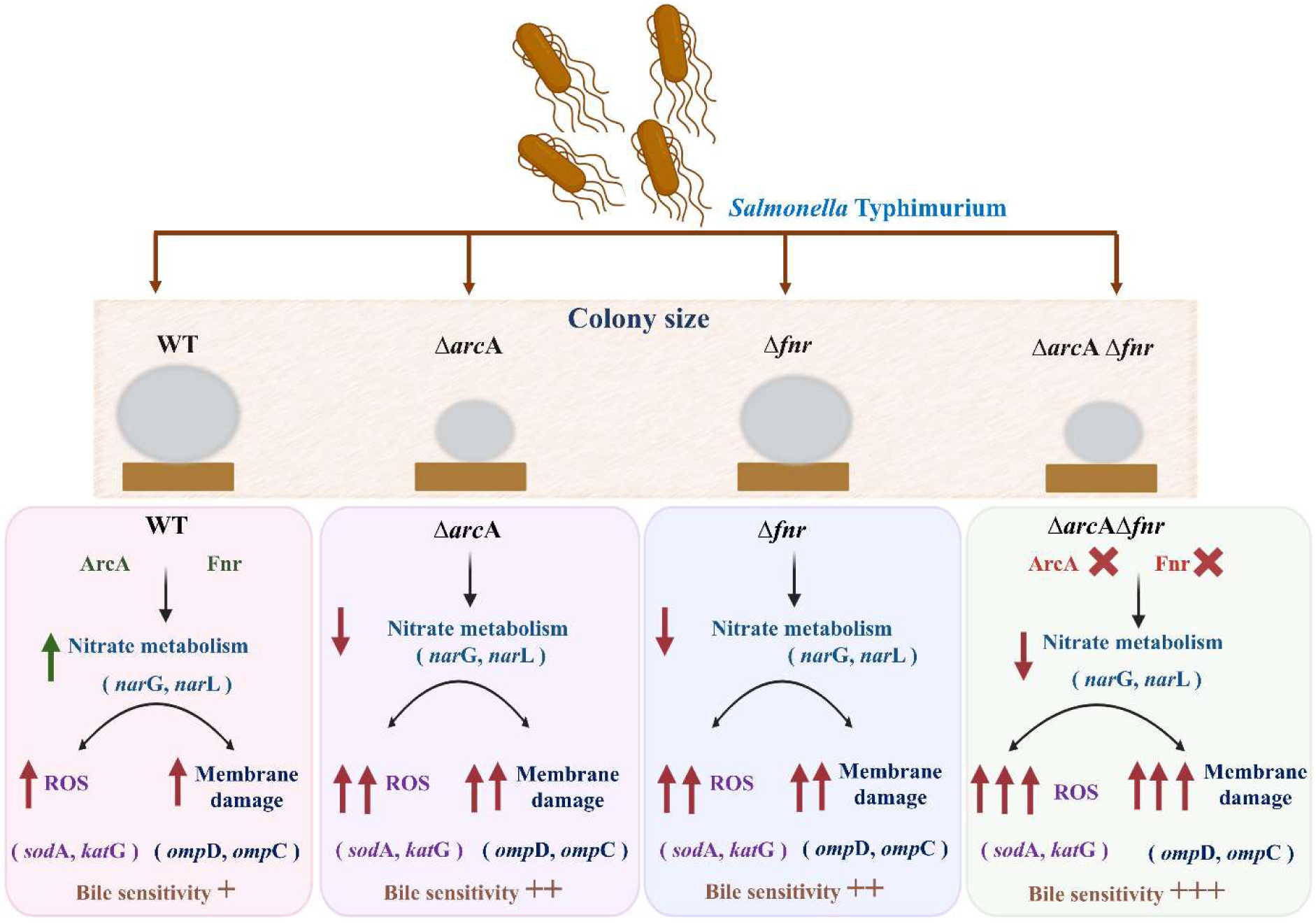
Schematic illustration depicting the roles of *S*. Typhimurium encoded ArcA and Fnr in mediating resistance to bile stress. *S*. Typhimurium encoded ArcA is important in determining the colony size and both ArcA and Fnr are essential for motility. The Δ*arc*AΔ*fnr* strain shows a significant decrease in colony size and impaired motility. Exposure to bile increases the expression of ArcA and Fnr and activation of the nitrate metabolism. This induction in nitrate metabolism is essential as the strain that lack this, such as, Δ*arc*A, Δ*fnr* and Δ*arc*AΔ*fnr* demonstrates high ROS and decreased membrane integrity. Overall, the co-regulation by ArcA and Fnr mediated nitrate metabolism is paramount for survival of *S*. Typhimurium in bile rich environment. Image was created using Biorender (Created in https://BioRender.com).

An earlier study in *E .coli* reported *fnr* dependent activation of *arc*A under anaerobic conditions, whereas under low oxygen conditions *fnr* exerts only a minor or negligible influence on *arc*A expression (Shalel-Levanon et al., 2005). *E. coli* encoded *fnr* activates transcription of *arc*A, resulting in an approximately fourfold increase in *arc*A expression in anaerobiosis. In a *fnr* deletion background there is no autoactivation of *arc*A, indicating that ArcA enhances its activation by Fnr. Also, Fnr activation of *arc*A transcription contributes to the coordination of genes associated with aerobic and anaerobic metabolism during environmental changes (Brown et al., 2022; Compan & Touati, 1994). Another study shows that in *E. coli*, Fnr directly activates transcription of *arc*A, and ArcA is required for oxidation of ubiquinol by nitrate reductases (NarG or Nap), linking *fnr* to *arc*A and electron transport processes (Constantinidou et al., 2006). Our findings also show that *arc*A and *fnr* work together to adapt to bile stress as both the single mutants show partial bile sensitivity (Fig. 1C). Time course expression analysis showed that the peak expression of *fnr* was induced post 3 h of bile exposure, whereas *arc*A showed a delayed induction at 5 h (Fig. 1A). It was also observed that the *fnr* mutant had a much lower level of *arc*A mRNA at 3 h, whereas the *arc*A mutant also showed lower amounts of *fnr* mRNA levels at 5 h (Fig. 1B) indicating a time-dependent regulatory interaction between *arc*A and *fnr*. The effects of *fnr* and *arc*A expression when either gene was deleted showed that they both regulate each other; however, the regulation of *arc*A by *fnr* is more pronounced. Importantly, both genes cooperate to enhance survival during bile stress.

All the four strains WT, Δ*fnr*, Δ*arc*A and Δ*arc*AΔ*fnr* exhibited no growth defect when grown in LB broth (Fig. 1C), however, their colony morphology on solid agar plates show distinct differences. WT and Δ*fnr* showed similar colony size on LB agar plate, whereas the Δ*arc*A and Δ*arc*AΔ*fnr* strains displayed smaller colony sizes (Fig. 2A-B). No difference was observed in the bacterial cell size among the four strains when cultured in either LB broth or LB agar plate (Fig. 2C-D). To our knowledge, this is the first time that the small colony phenotype has been seen in Δ*arc*A and Δ*arc*AΔ*fnr* strains. Cell motility, colony formation, and the diffusion of nutrients, oxygen, and metabolites have a significant impact on microbial growth (Chakraborty et al., 2025). Bacteria have the fastest growth rates in liquid food systems because they can move freely and diffuse effectively and growth rates may be higher under planktonic compared to surface conditions (Skandamis & Jeanson, 2015).

Flagellar biogenesis is governed by a transcriptional hierarchy consisting of Class I (*flh*DC), Class II (*fli*A), and Class III (*mot*A and *fli*L) genes, which together regulate flagellar assembly, motor function, and chemotaxis in *S.* Typhimurium (Chevance & Hughes, 2008; Ray et al., 2020). Previous reports in *S.* Typhimurium have shown that the *arc*A mutant was non motile and devoid of flagella yet retained virulence comparable to the WT strain (Evans et al., 2011). The phosphorylation site of ArcA was shown to be essential for motility, whereas its cognate sensor kinase ArcB was not required in *E. coli* (Kato et al., 2007). In *Vibrio cholerae*, deletion of *arc*A led to enhanced surface motility under aerobic conditions, by directly repressing the *flr*A expression, suggesting that *arc*A functions as a negative regulator of motility in this organism (Li et al., 2022). Deletion of *arc*B, but not *arc*A, significantly impaired motility in *Serratia marcescens* (Zhang et al., 2018). Transcript levels of *flh*D were significantly reduced in Δ*arc*A, whereas Δ*fnr* displayed lower transcripts of *flh*D*, fli*A and *fli*L as compared to WT, Δ*arc*AΔ*fnr* strain on the other hand, showed lower transcript of *flh*D, *fli*A, *fli*L and *mot*A as compared to WT strain, indicating impaired expression of Class I, II, and III flagellar genes (Fig. 3A). *Salmonella* exhibits two different types of flagella driven motility: swimming and swarming. In bacteria, swimming motility occurs in liquid environments or in media with low percentage agar (0.1–0.3%) using flagellar filaments. Swarming is a surface associated, multicellular behaviour observed in flagellated bacteria often observed in media with higher percentage of agar (Thakur et al., 2020; Wadhwa & Berg, 2022). In this study, deletion of *arc*A or *fnr* resulted in reduced swimming and swarming motility, whereas loss of both regulators caused a severe motility defect (Fig. 3B-D). The severe motility defect observed in the Δ*arc*AΔ*fnr* strain suggests that *arc*A and *fnr* act through partially overlapping pathways to regulate flagellar gene regulation, leading to severely impaired motility. Previous studies showed reduced motility of Δ*arc*A in *S*. Typhimurium (Evans et al., 2011) and Δ*fnr* in *S.* Typhimurium (Fink et al., 2007), however we found that the Δ*arc*AΔ*fnr* strain exhibited an even more pronounced defect.

Bile has been shown to directly damage the outer membrane and affect porins and efflux pumps in *S.* Typhimurium. The bile sensitive Δ*csp*E mutant displays higher expression of multiple porins, such as OmpC, OmpF, and OmpD, which may lead to greater bile sensitivity (Ray et al., 2019). *E. coli* encoded OmpC excludes bile salts due to smaller pore size and contributes to cell survival. It has been previously demonstrated that OmpC contributes to adaptation within the intestinal environment and is preferentially expressed in the presence of bile salts (Doranga & Conway, 2023). Thus, the higher ompC expression in the Δ*arc*A and Δ*arc*AΔ*fnr* (Fig. 4A) could be a compensatory response to increased envelope damage caused by bile. Compared to the untreated control, bile salt treated Δ*arc*AΔ*fnr* strain showed noticeably greater intracellular accumulation of bisBenzimide H 33258 (Fig. S7A-B) and NPN dyes as compared to the single mutants (Fig. 4B-C). This increase is possibly due to the bile treated Δ*arc*AΔ*fnr* strain’s higher porin levels, which may have promoted bile influx and enhanced cellular permeability. Increased *omp*C transcript levels reflect altered outer membrane composition, while elevated NPN uptake indicates compromised outer membrane integrity. Whether increased porin expression directly contributes to membrane damage or reflects a parallel envelope stress response remains to be determined.

Bile exposure increases the generation of intracellular ROS in *S*. Typhimurium (Singh et al., 2023). Also, *arc*A and *fnr* mutant fail to survive in macrophages due to their inability to detoxify the ROS that is necessary for bacterial survival during infection (Fink et al., 2007; Pardo-Esté et al., 2018). In the present study, bile treatment resulted in a marked increase in intracellular ROS amounts, with the highest ROS amounts observed in the Δ*arc*AΔ*fnr* strain (Fig. 5B). This was accompanied by a significant upregulation of *sod*A and *kat*G (Fig. 5A), which encode superoxide dismutase and catalase, respectively. The severe ROS burden observed in the Δ*arc*AΔ*fnr* strain is consistent with an inability to survive under bile induced stress (Fig. 1C). Transient increase in nitrate metabolism may help cells cope with bile stress by utilizing nitrate as an alternative electron acceptor, thereby reducing reliance on oxidative metabolism and limiting ROS accumulation (Fig. 6B). Bile-sensitive Δ*arc*A, Δ*fnr*, and Δ*arc*AΔ*fnr* strains that induced high ROS were unable to produce nitrite, suggesting compromised nitrate metabolism (Fig. 6B). Nitrate metabolism is functionally impaired in the mutants because nitrite is a direct byproduct of nitrate reduction via the NarGHIJ and NarXL system. Previous multi-omics research in *E. coli* has also shown that *arc*A and *fnr* are activated under nitrate conditions (Toya et al., 2012). Whether *arc*A also regulates the nitrate metabolism upon bile exposure along with *fnr* in *S*. Typhimurium was unknown. While *fnr* is known to regulate nitrate metabolism under bile stress, our results suggest a novel role for *arc*A in nitrate metabolism indicating that these global regulators collectively contribute to metabolic adaptation under bile stress. Nitrate pretreatment significantly decreased intracellular ROS levels in all strains (Fig. 7A). Pretreating WT, Δ*arc*A, Δ*fnr*, and Δ*arc*AΔ*fnr* strains with sodium nitrate also reduced NPN fluorescence (Fig. 7B) (Fig. S9A), and intracellular Hoechst dye accumulation (Fig. S9B), corresponding to decrease in outer membrane permeability. These findings suggest that by providing nitrate as an alternative terminal electron acceptor and reducing redox pressure, partially compensates for the impaired nitrate metabolism in the deletion mutants during bile treatment. Significantly, pretreatment with antioxidants like GSH and NAC partially rescued the bile sensitivity (Fig. S10 A-B), and decreased intracellular ROS levels in Δ*arc*AΔ*fnr* strains (Fig. 7D). However, it did not restore nitrite production (Fig. 7E), suggesting that scavenging ROS is insufficient to replicate the bile adaptive phenotype similar to WT in these deletion mutants. This result indicates that, *arc*A and *fnr* dependent nitrate respiration is the major contributor to redox balancing during bile stress.

An important outcome of this study is the rescue of bile sensitivity in Δ*arc*A, Δ*fnr*, and Δ*arc*AΔ*fnr* strains by terminal electron acceptors. Among the tested electron acceptors, nitrate represents the most energetically favourable electron acceptor available to *S.* Typhimurium. The nitrite/nitrate redox couple has a standard redox potential (E° ∼ +433 mV), allowing efficient electron withdrawal from the respiratory chain (Rivera-Chávez & Bäumler, 2015). Fumarate reductases play a crucial role in maintaining cellular redox homeostasis by reoxidizing reduced electron carriers such as FADH₂ and NADH under oxygen limited conditions (Kim et al., 2018). Anaerobic respiration can also be supported by dimethyl sulfoxide (DMSO) that enhance gut colonization of *Salmonella* (Cruz et al., 2023). The exogenous addition of both sodium fumarate and DMSO rescued the bile sensitivity of Δ*arc*A, Δ*fnr*, and Δ*arc*AΔ*fnr* strains, however exogenous addition of sodium nitrite failed to rescue the bile sensitivity (Fig. S8). Nitrite possesses antimicrobial activity due to its ability to generate nitric oxide under acidic conditions, inactivate iron-sulfur proteins and inhibit bacterial respiration (Mühlig et al., 2014). It is likely that exogenous addition of sodium nitrite does not relieve bile stress due to its cytotoxic effects.

ArcA and Fnr work together to co regulate a large number of genes involved in energy metabolism and stress adaptation, as well as to coordinate metabolic transitions in response to oxygen availability. They coregulate many genes, including those encoding cytochrome c oxidase (cyoABCDE), tricarboxylate transport, ethanolamine utilization, L-lactate transport and metabolism, the aerotaxis sensor receptor (aer) and nitrite reductase (nrfAB) (Evans et al., 2011). Overall, this study indicates that disruption of ArcA and Fnr regulatory pathway lowers nitrate metabolism, resulting in higher ROS and membrane damage, which undermines bacterial survival upon bile exposure. Our results show a previously unidentified function of *arc*A in the regulation of nitrate metabolism, under bile stress. Also, nitrate metabolism functions upstream of redox homeostasis during bile stress adaptation, wherein *arc*A and *fnr* govern nitrate metabolism that negatively influences intracellular ROS accumulation. These results demonstrate a collaborative function of ArcA and Fnr in enhancing survival of *S.* Typhimurium to bile stress.

## Materials and methods

### Bacterial strains and growth conditions

*Salmonella* Typhimurium 14028s was used as the WT strain. Table S1 lists the plasmids and bacterial strains used in this investigation. Cultures were cultivated at 37°C with constant aeration at 180 rpm in Luria Bertani (LB) broth that contained 10 g/litre NaCl (Merck, Darmstadt, Germany), 10 g/litre tryptone (Himedia Laboratories, Mumbai, India), and 5 g/litre yeast extract (Himedia Laboratories). Every experiment used overnight grown single colony cultures of WT, Δ*arc*A, Δ*fnr* and Δ*arc*AΔ*fnr* strain as the pre inoculum. pKD46 carrying strains were cultivated at 30°C with continuous aeration at 180 rpm. Bile salts containing cholic acid-deoxycholic acid sodium salt mixture (Sigma-Aldrich, St. Louis, MO, USA) were used at concentrations of 1-5% (w/v) or as mentioned otherwise. LB containing antibiotics were used at the following concentrations: kanamycin, 50 μg/ml and ampicillin, 100 μg/ml. pQE60^ampR^ was used for overexpression of *arc*A and *fnr*.

### Preparation of gene deletion strains

The Δ*csp*E (Ray et al., 2019) and Δ*fnr* (Singh et al., 2024) strains were used from an earlier study. The Δ*arc*A single deletion and Δ*arc*AΔ*fnr* double deletion strain were generated using the one step gene disruption strategy (Datsenko & Wanner, 2000). Table S2 lists the primers (Sigma, Bangalore, India) used to create the Δ*arc*A strain. The *kan* from pKD4 with 40 nucleotide *arc*A flanking regions was amplified using these primers. After purifying and electroporating the resultant PCR product into WT cells carrying pKD46, transformed cells were chosen for gene deletion on Kanamycin (50 μg/ml) (VWR Chemicals, Sanborn, NY, USA) plates. By using pcp20 for transformation, the kanamycin cassette was eliminated. The double deletion strain Δ*arc*AΔ*fnr* were generated by amplifying the region flanking *fnr* and electroporating the amplicon in Δ*arc*A cells harbouring pKD46. Using pCP20, antibiotic resistance cassettes were eliminated. All gene deletions were validated by PCR amplification using primers listed in TableS4.

### Stress assays

All the overnight grown cultures were normalized O.D._600_ of 2. The experiments were performed in 5 ml LB followed by addition of bile to make up the final concentration of 3% v/v. For the kinetics experiment bile was added in beginning and *A* at 600 nm was recorded using a Microplate reader (Tecan, Grodig, Austria) in a clear flat bottom 96 well culture plate (Tarsons, Kolkata, India) every 3 h. Appropriate dilutions were plated on LB agar plates and C.F.U. was recorded after overnight incubation at 37°C. For complementation experiments, the overnight grown cultures were normalized to O.D._600_ of 2 and bile was added and incubated for 6 h at 37°C (Ray et al., 2019; Singh et al., 2024). The *A* at 600 nm was recorded at 6 h and appropriate dilutions were plated to record the C.F.U. For H_2_O_2_ assay the cells were adjusted to O.D._600_ of 2 and were inoculated in 50 ml LB with 0.5 mM H_2_O_2_, 0.75 mM H_2_O_2_, 1mM H_2_O_2_. The cells were incubated for 6 h at 37°C and *A* at 600 nm was recorded using a Microplate reader, as mentioned above.

### Colony morphology estimation

Overnight grown cultures were normalized to O.D._600_ of 2 and appropriate dilutions were plated on LB agar plates. These plates were allowed to be incubated for 12 to 48 h, at 37°C. Images for the colony morphology were obtained by a stereo microscope (*Nikon* SMZ745T MICAPS HPS3CMOS, Japan) at 20x (Chakraborty et al., 2025).

### Atomic Force microscopy

Overnight cultures of WT, Δ*arc*A, Δ*fnr* and Δ*arc*AΔ*fnr* strains were normalized to O.D._600_ of 2 and incubated at 37 °C for 5 h. The cells were washed thrice with freshly autoclaved MQ water and the O.D._600_ was adjusted to 0.1. The normalized cultures (70 μl) were drop casted on clean cover slips and allowed to air dry overnight. After washing three times with freshly autoclaved MQ water, they were again allowed to dry completely. Once the samples were fully dried, AFM imaging (Park Systems, South Korea) was performed and cellular lengths were measured using the image processing software XEI (Chakraborty et al., 2025).

### Motility assays

The overnight cultures were normalized to O.D._600_ of 1. Swimming agar (0.3 % soft agar) was prepared freshly and 20 ml of soft agar was poured in the petri plates at room temperature. The plates were inoculated with 3 μl of the normalized culture at the centre of the plate and was incubated at 37°C. The images were acquired with ImageQuant LAS4000 (GE Healthcare) at 8 h and 16 h. For swarming motility, 0.5 % freshly prepared nutrient agar with 0.5 % glucose was poured into petri plates. The plates were allowed to dry at room temperature. 5 μl of the normalized culture was spotted in the middle of the plate and was incubated at 30°C. The images were captured at 8 h and 16 h. The diameters were estimated using ImageJ 1.x. For RNA isolation, each biological replicate represents RNA isolated from an independent swimming plate inoculated from separate overnight cultures (Ray et al., 2020).

### NPN dye uptake assay

Pure and complemented strains were grown overnight and adjusted to O.D._600_ of 2. The cells were inoculated in 5 ml LB with and without 3% bile for 6 h at 37 °C, 180 r.p.m. The cells were washed thrice with 1x PBS and adjusted to O.D._600_ of 0.5 and incubated with 10 μM of N-phenyl-1-naphthylamine (NPN) and kept in dark for 10 min and analysed by flow cytometry (BC Cytoflex LX, USA). Data were analysed using FlowJo software (version 10) using a single gate applied uniformly across all samples and conditions. For the NaNO_3_ pretreatment the cells were treated with 25 mM sodium nitrate, incubated at 37 °C, with aeration at 180 r.p.m., grown to O.D.600 of 0.2 and then treated with bile (final concentration 3% v/v) for 6 h. The samples were stained with 10 μM of N-phenyl-1-naphthylamine (NPN) and proceeded towards analysis by flow cytometry (Chakraborty et al., 2025).

### ROS estimation

Overnight grown cultures were normalized to an O.D._600_ of 2 and inoculated into 5 ml LB. Cultures were grown till O.D._600_ of 0.2, followed by treatment with 3 % bile at 37 °C, 180 rpm for 5.5 h or the time mentioned. Cells were washed thrice with 1× PBS and incubated with 20 μM 2′,7′-Dichlorofluorescin Diacetate (DCFDA; Sigma-Aldrich) at 37 °C for 30 minutes. After incubation, excess dye was removed by washing, and 200 μl of the cell suspension was transferred to a 96 well plate. Fluorescence was measured using an Infinite 200 Pro plate reader (Tecan, Austria GmbH) at excitation/emission wavelengths of 485/535 nm. Fluorescence values were normalized to the O.D._600_ of each sample. For the ROS estimation upon sodium nitrate pretreatment, the normalized culture was incubated at 37 °C, 180 rpm, till O.D._600_ of 0.2 with 25 mM of sodium nitrate and then treated with 3 % bile for 5.5 h (Singh et al., 2023).

### Griess assay to measure intracellular nitrite concentration

Intracellular nitrite was estimated using the Griess assay (Singh et al., 2024). Overnight cultures of WT, Δ*arc*A, Δ*fnr* and Δ*arc*AΔ*fnr* was adjusted to O.D._600_ of 0.2 and then treated with bile for the time points mentioned at 37 °C, 180 r.p.m. The cells were washed twice with 1x PBS and sonicated (QSonica Q700). Post cell lysis, the cells were centrifuged at 13000 g for 15 min. 150 μl of supernatant was incubated with 100 μl of Griess reagent at room temperature in dark. O.D._570_ was recorded using a microplate reader (Tecan, Austria Gmbh).

### Assessment of bile tolerance under sodium nitrate supplementation

Overnight cultures of WT, Δ*arc*A, Δ*fnr* and Δ*arc*AΔ*fnr* was adjusted to O.D._600_ of 2. The cells were inoculated in 5 ml LB with and without supplementation with sodium nitrate. The cells were allowed to grow till O.D._600_ of 0.2 and then treated with bile for 6 h at 37 °C, 180 rpm. O.D._600_ was recorded using a microplate reader and appropriate dilutions were plated in LB agar plate to calculate the CFUs (Singh et al., 2024).

## Statistical analysis

All statistical analyses in this work were performed in GraphPad Prism 8 using ANOVA and Turkey’s multiple comparison test wherever applicable. Data were represented as mean ± S.D., where *p ≤ 0.05, **p ≤ 0.01, ***p ≤ 0.001, and ****p ≤ 0.0001.

