## Supplemental Information for "Coregulation by *arc*A and *fnr* protects *Salmonella* Typhimurium from bile stress by maintaining redox homeostasis and membrane integrity"

**Supplementary Tables**

**Table S1: List of strains and plasmids**

| Strain | Genotype/ features | Reference |
|---|---|---|
| *S*. Typhimurium 14028s (WT) | Wild-type, parent of all *Salmonella* strains | (Singh et al., 2024) |
| Δ*csp*E | 14028s; *csp*E::FRT | (Singh et al., 2024) |
| Δ*arc*A | 14028s; *arc*A::FRT | This study |
| Δ*fnr* | 14028s; *fnr*::FRT | (Singh et al., 2024) |
| Δ*arc*AΔ*fnr* | 14028s; *arc*A:: FRT , *fnr*:: FRT | This Study |
| WT/VA | 14028s; pQE60; $Amp^r$ | This Study |
| Δ*arc*A/VA | 14028s; pQE60; $Amp^r$ | This Study |
| Δ*fnr*/VA | 14028s; pQE60; $Amp^r$ | This Study |
| Δ*arc*AΔ*fnr*/VA | 14028s; pQE60; $Amp^r$ | This Study |
| WT/ p*arc*A | 14028s; pQE60; $Amp^r$ | This Study |
| Δ*arc*A/ p*arc*A | 14028s; pQE60; $Amp^r$ | This Study |
| WT/ p*fnr* | 14028s; pQE60; $Amp^r$ | This Study |
| Δ*fnr*/ p*fnr* | 14028s; pQE60; $Amp^r$ | This Study |
| Δ*arc*AΔ*fnr*/ p*arc*A | 14028s; pQE60; $Amp^r$ | This Study |
| Δ*arc*AΔ*fnr*/ p*fnr* | 14028s; pQE60; $Amp^r$ | This Study |
| pQE60 | A 3431 bp bacterial expression plasmid. Contains T5 promoter and confers ampicillin resistance | (Singh et al., 2024) |
| pKD46 | Temperature sensitive λ-Red recombinase expression plasmid; $Amp^r$ | (Singh et al., 2024) |
| pKD4 | Kanamycin resistance (*kan*) | (Singh et al., 2024) |
| pCP20 | Temperature sensitive FLP recombinase expression plasmid; $Amp^r$ | (Singh et al., 2024) |
| p*arc*A | pQE60 with *arc*A cloned between BamHI and HindIII | This Study |
| p*fnr* | pQE60 with *fnr* cloned between BamHI and HindIII | (Singh et al., 2024) |

**Table S2: List of primers used for gene knockout preparation**

| Gene | Sequence 5`- 3` |
|---|---|
| *arc*A FP | ACTTCCTGTTTCGATTTAGTTGGCAATTTAGGTAGCAAACGTGTAGGCTGGAGCTGCTTC |
| *arc*A RP | TTAATCCTGCAGGTCGCCGCAGAAGCGATAACCTTCGGAATATCCTCCTTAGTTCCT |

**Table S3: List of primers used for gene knockout confirmation**

| Gene | Sequence 5`- 3` |
|---|---|
| *arc*A FP | AAAAACGGTCTCCTGTTAGC |
| *arc*A RP | TAGTTTATACTGCTCGCCG |
| *fnr* FP | TCAATTACGGCTTGAGCAG |
| *fnr* RP | CATCAATGGTTTAGCTGACG |

**Table S4: List of primers used for cloning**

| Gene | Sequence 5`- 3` |
|---|---|
| *arc*A FP | CGCGGATCCATGCAGACCCCGCACATTCT |
| *arc*A RP | CCCAAGCTTTTAATCCTGCAGGTCGCCGC |

**Table S5: List of primers used for qRT-PCR**

| Gene | Sequence 5`- 3` |
|---|---|
| *fnr* FP | TTAACGAGCATGAGCTTGAT |
| *fnr* RP | TGTAGCTCTTAATCGTTCCG |
| *arc*A FP | AAAAACGGTCTCCTGTTAGC |
| *arc*A RP | TAGTTTATACTGCTCGCCG |
| *flh*D FP | CATTTGACCTTTTTGCTTCT |
| *flh*D RP | GTTTTAGCAACTCGGATGTA |
| *mot*A FP | AAAAATCACACTGCTGTCTA |
| *mot*A RP | CCAACTCAATAAACGATGGA |
| *fli*A FP | ACAGCCAACTTTTCTCTTAC |
| *fli*A RP | AGTTGATGTAACGGGTTTTC |
| *fli*L FP | CTGGCAAACAAAAACTGATT |
| *fli*L RP | TTACCGCAGAATAAAAGCTG |
| *omp*C FP | TGAAATACGATGCGAACAAC |

| | |
|---|---|
| *omp*C RP | TACTGAGCAACCACTTCAAA |
| *sod*A FP | ATAAGCAGACGATGGAGATT |
| *sod*A RP | TAGTAATCAGTTCTTCAACCG |
| *kat*G FP | GAGGTAAAATACACGGCGA |
| *kat*G RP | GGTTCATCACTTTCACCCAT |
| *nar*G FP | GAAAGTTTGCTGGTGTATCG |
| *nar*G RP | CAGCATCAACAGATTATCGC |
| *nar*L FP | TGGTTGGCGGTATTTTACTG |
| *nar*L RP | TCATTTCACTCCCGTCCATA |

**Table S6: Intra strain relative expression levels of *arc*A, *fnr*, *gmk* and *gyr*B in *S.* Typhimurium D23580 strain (SalCom V2.0)**

| **Conditions** | ***arc*A** | ***fnr*** | ***gmk*** | ***gyr*B** |
|---|---|---|---|---|
| EEP | 1 | 1 | 1 | 1 |
| MEP | 0.77 | 1.05 | 0.79 | 0.8 |
| LEP | 0.93 | 0.93 | 0.55 | 0.57 |
| ESP | 1.46 | 0.64 | 0.44 | 0.44 |
| LSP | 0.27 | 0.32 | 0.1 | 0.08 |
| MEP | 1 | 1 | 1 | 1 |
| NaCl shock | 1.12 | 0.73 | 0.46 | 0.43 |
| **Bile shock** | **1.34** | **1.49** | **1.77** | **1.55** |
| Low Fe2+ shock | 0.9 | 0.84 | 0.74 | 0.54 |
| Anaerobic shock | 1.66 | 0.49 | 0.69 | 0.3 |
| Anaerobic growth | 1 | 1 | 1 | 1 |
| Oxygen shock | 0.72 | 1.65 | 1.02 | 1.68 |
| InSPI2 | 1 | 1 | 1 | 1 |
| Peroxide shock (InSPI2) | 0.97 | 0.86 | 0.49 | 0.26 |
| Nitric Oxide shock (InSPI2) | 1.87 | 0.78 | 0.95 | 0.6 |
| NonSPI2 | 1 | 1 | 1 | 1 |
| InSPI2 | 0.67 | 0.75 | 0.62 | 0.63 |
| ESP | 1 | 1 | 1 | 1 |
| Macrophage | 1.14 | 1.6 | 2.29 | 1.32 |

**Supplementary Fig. 1**

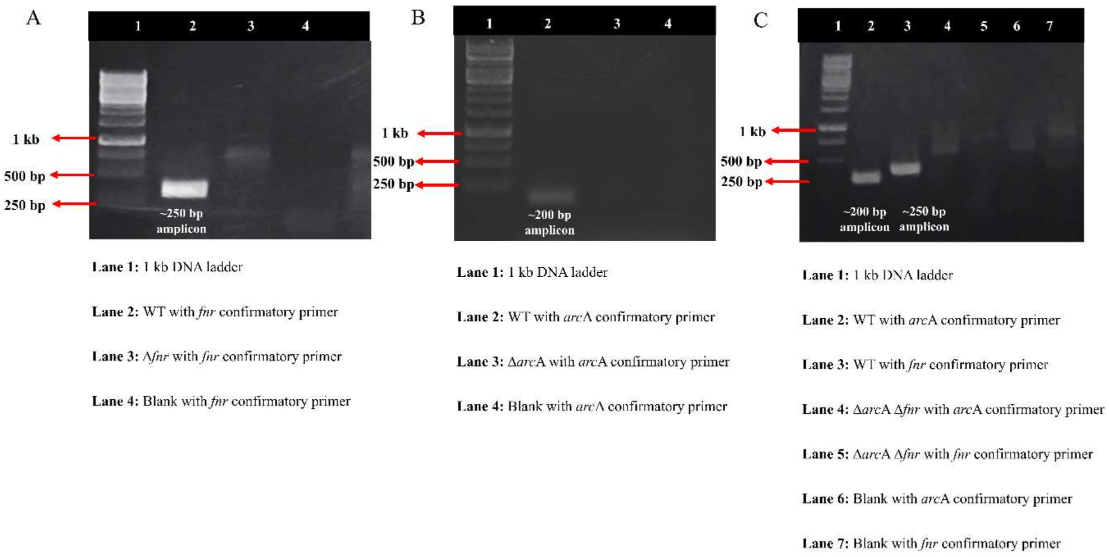


**Fig. S1 PCR confirmation for the Δ*arc*A, Δ*fnr* and Δ*arc*AΔ*fnr* strains.** A colony PCR was performed to confirm the strains. DNA ladder of 1 Kb was used to determine presence of amplicons. WT 14028s was used as positive control. Gene knockouts of Δ*arc*A, Δ*fnr* and Δ*arc*AΔ*fnr* were confirmed by running on 1% agarose gel.

### Supplementary Fig. 2

**(A)**

| WT | | | WT+ 3% Bile | | | |
|---|---|---|---|---|---|---|
| 4.22159 | 3.47305 | 4.68985 | 2.68181 | 2.47615 | 1.6529 | *dms* B |
| 3.61347 | 2.71526 | 4.0415 | 1.34621 | 1.16948 | 0.91593 | *dms* C |
| 3.74714 | 4.30022 | 4.8963 | 2.16335 | 2.41341 | 2.05344 | *pdu* F |
| 4.12664 | 5.01604 | 5.83564 | 0.41333 | 0.09148 | -0.66903 | *pdu* A |
| 2.21237 | 2.76679 | 3.99998 | -0.72418 | 0.09148 | -1.406 | *pdu* J |
| 0.83386 | 1.32621 | 2.45654 | -1.46114 | -3.07844 | -2.99096 | *pdu* K |
| 4.89851 | 5.00536 | 4.79069 | 1.70878 | 1.77954 | 0.70948 | *eut* M |
| 1.01443 | 0.63434 | 0.36138 | -0.72418 | -0.27109 | -0.66903 | *eut* P |
| 0.6274 | 0.84079 | 0.36138 | -0.72418 | -0.75652 | -0.66903 | *eut* A |
| 0.83386 | 1.02136 | 0.60239 | -0.72418 | -0.27109 | -1.406 | *eut* B |

**(B)**

| **WT** | | | **WT + 3% Bile** | | | |
|---|---|---|---|---|---|---|
| 0.39 | 0.39 | -0.29 | 1.91 | 1.31 | 2.68 | ***arc* A** |
| 2.28 | 2.61 | 2.33 | 2.38 | 1.97 | 2.29 | ***fnr*** |

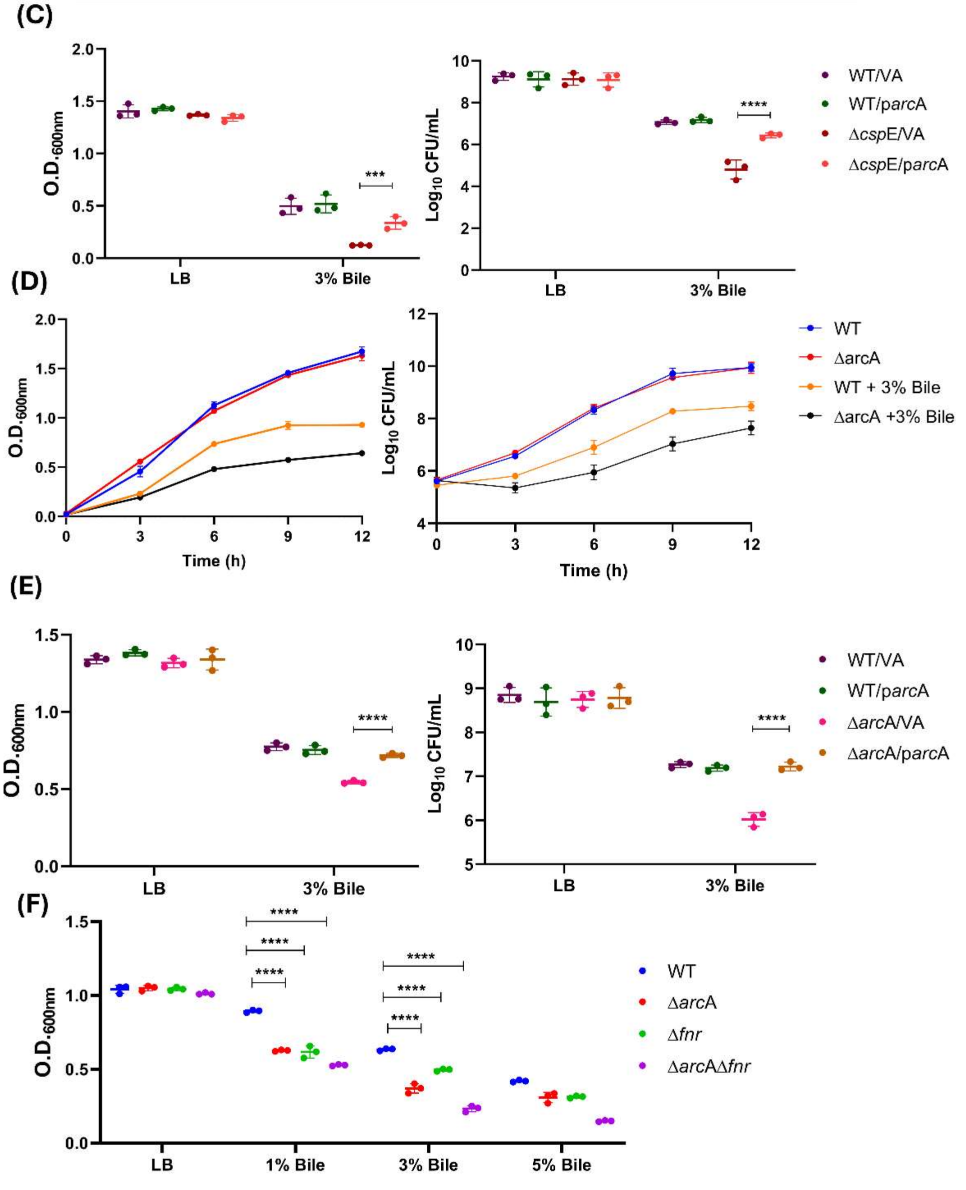

**Fig. S2 Bile treatment modulates *arc*A and *fnr* expression to promote bacterial viability.** (A) Heat map depicting differential expression of genes involved in anaerobic metabolism between bile treated and untreated WT strain (n=3) with >1 Log2 fold change. (B) Heat map depicting the differential expression values of *arc*A and *fnr* in WT upon treatment with bile (n=3) with >1 Log2 fold change. (C) Overexpression of *arc*A in Δ*csp*E. O.D. was recorded post bile treatment for 6 h and C.F.U. was performed after plating on LB plate with proper dilutions. (D) The growth of Δ*arc*A and WT strain with and without bile treatment. O.D. was plotted at appropriate time interval and C.F.U. was recorded after plating with proper dilutions at respective time interval. (E) Complementation with p*arc*A in *arc*A with and without the presence of 3% bile. (F) Indicated strains were treated with different amount of bile. The data are presented as mean ± SD and is representative of 3 independent experiments. $P$ values were measured by two-way ANOVA using Turkey's multiple comparison test. **$P$< 0.01, ***$P$<0.001, ****$P$ < 0.0001.

**Supplementary Fig. 3**

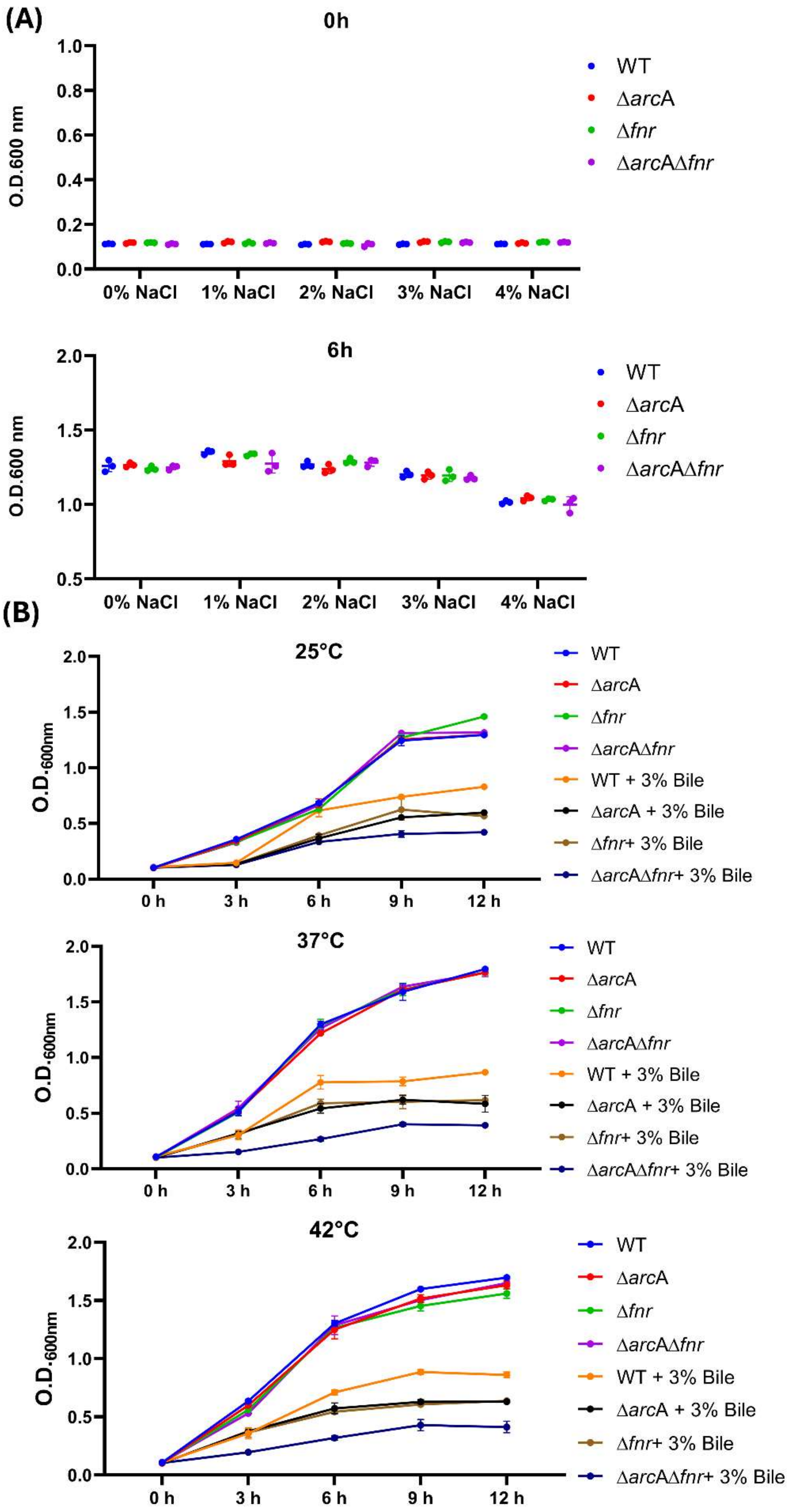

**Fig. S3 The growth of Δ*arc*A, Δ*fnr*, and Δ*arc*AΔ*fnr* strains remains unaffected by osmotic stress, and temperature variations.** O.D. was measured after treatment with (A) NaCl in a dose dependent manner at 0 h and 6 h and (B) growth in temperature at 25° C, 37° C and 42° C. Data is representative of 3 individual experiments.

**Supplementary Fig. 4**

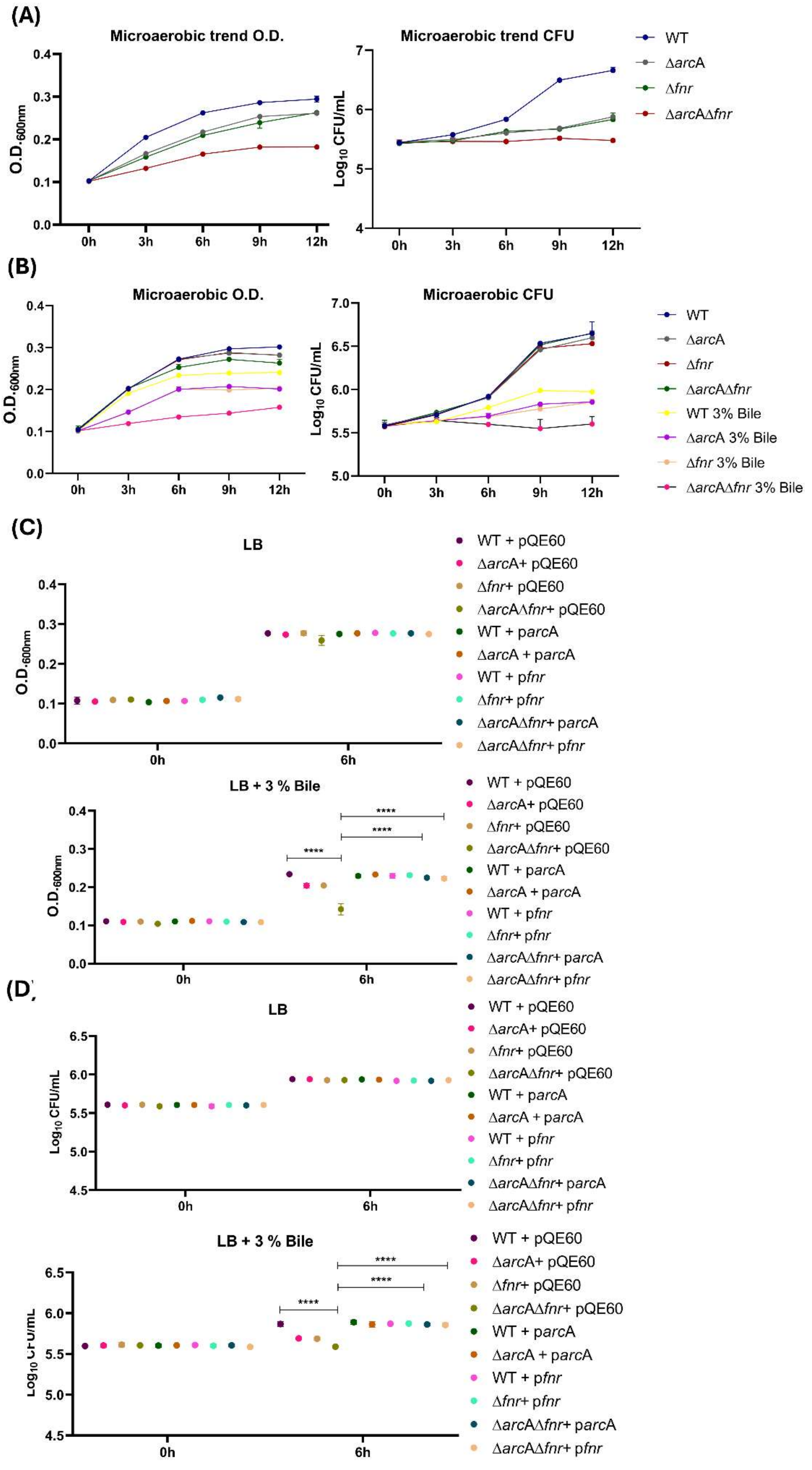

**Fig. S4 Under microaerobic stress the Δ*arc*AΔ*fnr* strain displays increased sensitivity to bile stress as compared to WT and single mutants Δ*arc*A and Δ*fnr*.** (A) A time kinetics for all the strains under microaerobic stress. (B) The growth of WT, Δ*arc*A, Δ*fnr* and Δ*arc*AΔ*fnr* strains with standardized volume of culture upon 3% bile treatment was performed in a time dependent manner. (C) Complementation of Δ*arc*A with p*arc*A, Δ*fnr* with p*fnr* and Δ*arc*AΔ*fnr* with p*arc*A and p*fnr* with 3% bile. CFU were counted after plating cells with appropriate dilutions. Data are presented as mean ± SD and is representative of 3 independent experiments. *P* values were measured by two-way ANOVA using Turkey's multiple comparison test. *****P* < 0.0001.

**Supplementary Fig. 5**

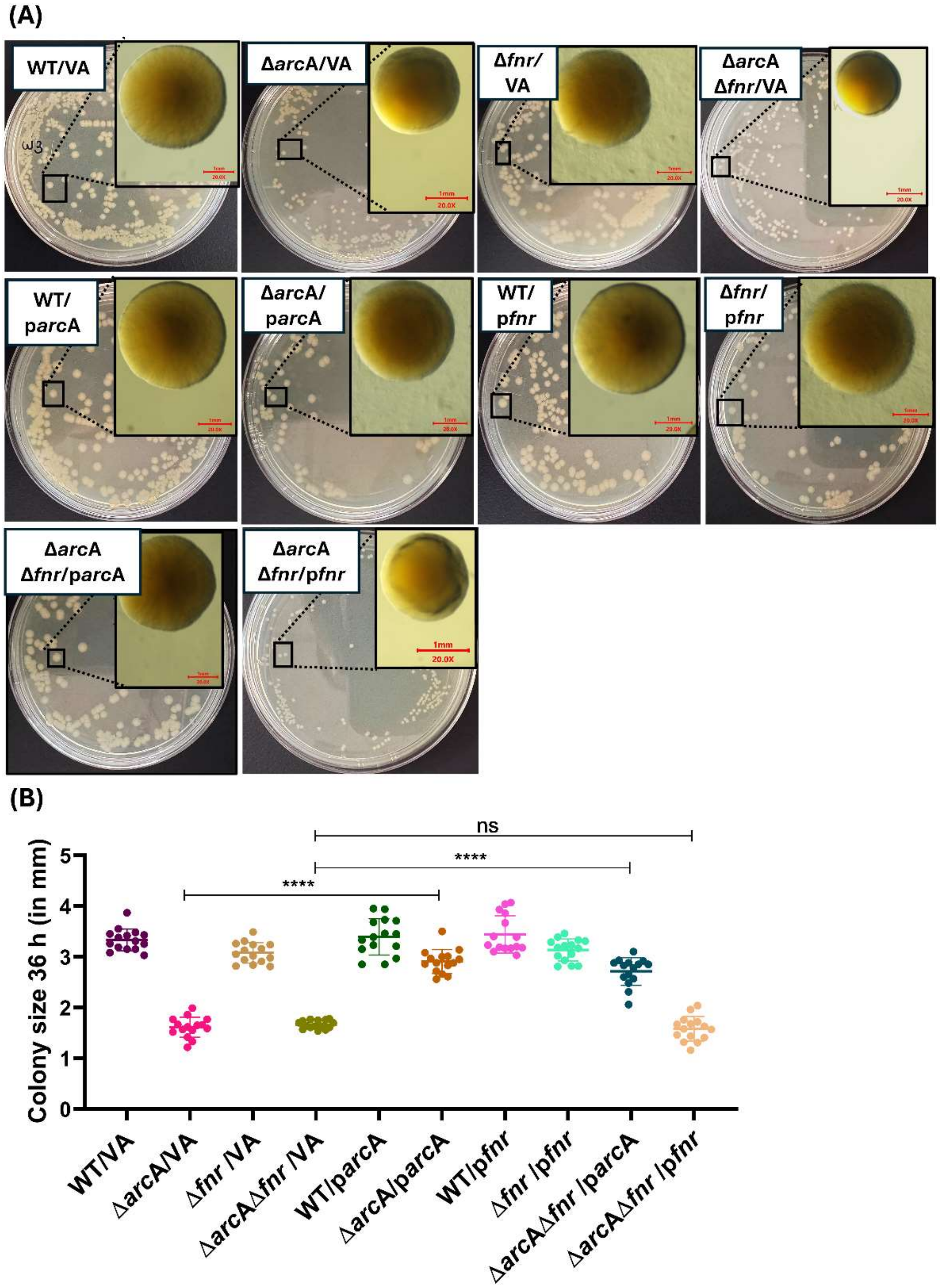


**Fig. S5 Complementation with p*arc*A rescues the reduced colony size of the Δ*arc*AΔ*fnr* strain.** (A) Depiction of colony morphology for the complemented strains on LB plates at 36 h and, (B) quantitation of the diameter for colonies at 36 h. The data is representative of 3 individual experiments and presented as mean ± SD. P values were measured by two-way ANOVA using Turkey's multiple comparison test. ****P < 0.0001.

### Supplementary Fig. 6

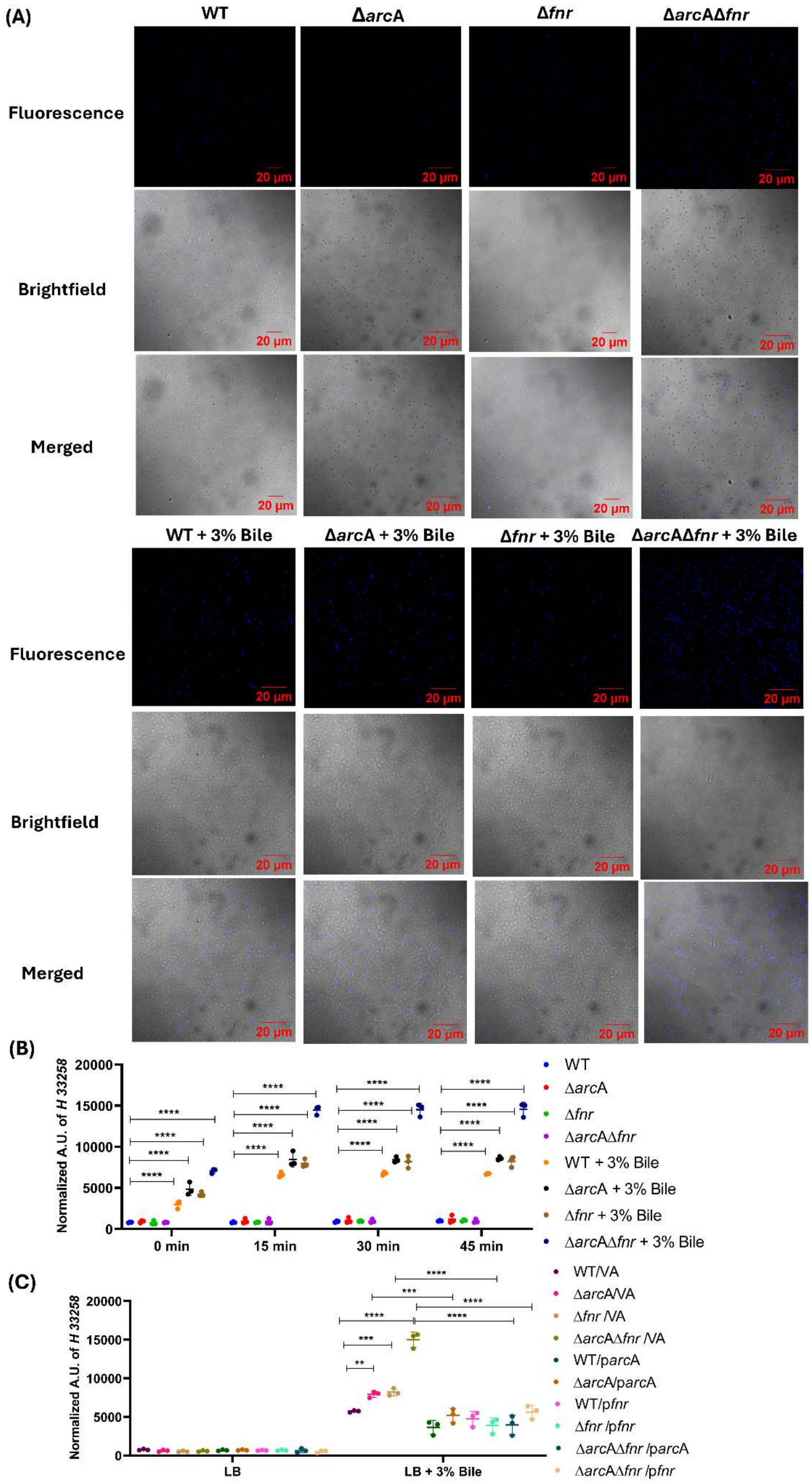

**Fig. S6 The Δ*arc*AΔ*fnr* strain exhibits increased permeability upon bile exposure as compared to Δ*arc*A, Δ*fnr* and WT.** (A) Representative fluorescence images of WT, Δ*arc*A, Δ*fnr* and Δ*arc*AΔ*fnr* post 30 min incubation with bisbenzimide H 33258 after addition of 3% bile for 5 h. (B) Quantification of bisbenzimide H 33258 accumulation in a time dependent manner in WT, Δ*arc*A, Δ*fnr* and Δ*arc*AΔ*fnr* cells after bile exposure. (C) Complementation with p*arc*A and p*fnr* rescued the increased permeability of bisbenzimide H 33258. Data is representative of 3 individual experiments and presented as mean ± SD. $P$ values were measured by two-way ANOVA using Turkey's multiple comparison test. *$P < 0.1$, **$P < 0.01$, ***$P < 0.001$, ****$P < 0.0001$.

**Supplementary Fig. 7**

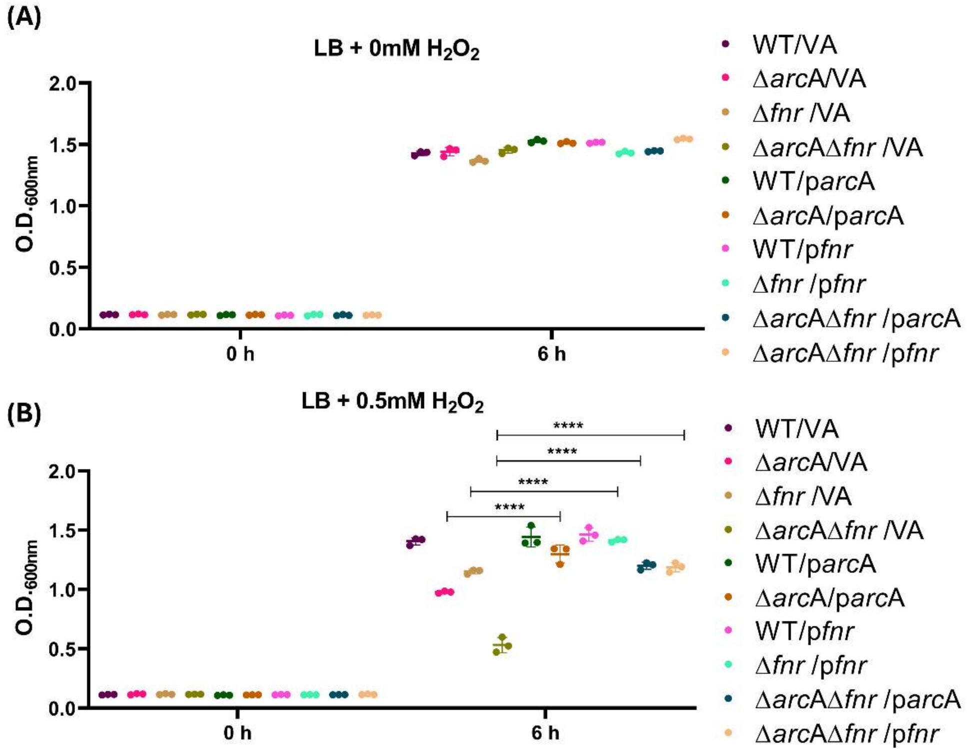


**Fig. S7 Complementation with p*arc*A and p*fnr* rescues the hydrogen peroxide sensitive phenotype of Δ*arc*A, Δ*fnr* and Δ*arc*AΔ*fnr*.** O.D. was measured post treatment with (A) 0mM $H_2O_2$ and (B) 0.5mM $H_2O_2$. Data is representative of 3 individual experiments and presented as mean ± SD. *P* values were measured by two-way ANOVA using Turkey's multiple comparison test. ****$P < 0.0001$.

**Supplementary Fig. 8**

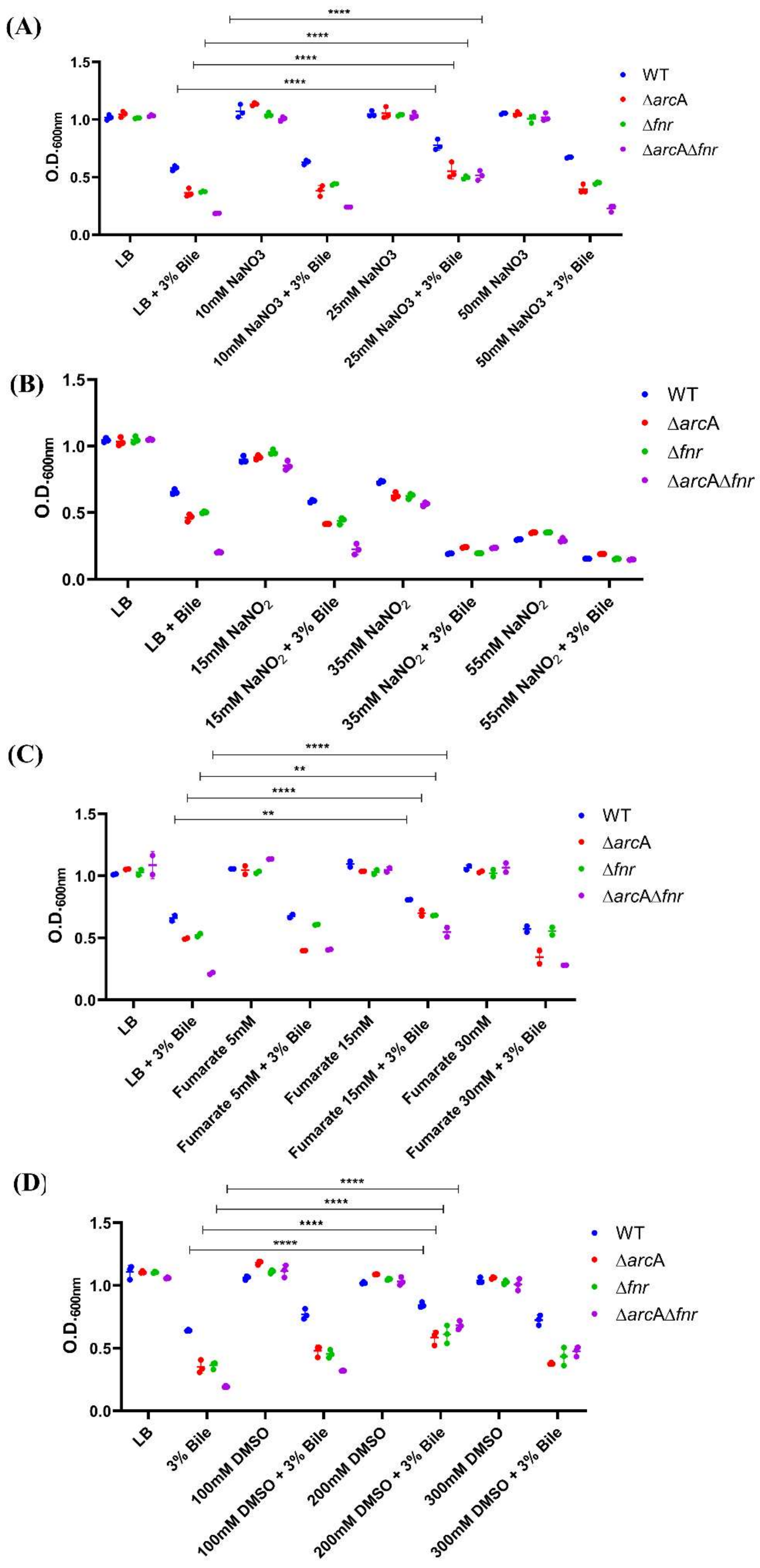

**Fig. S8 Pretreatment with sodium nitrate, sodium fumarate and DMSO rescues the bile sensitive phenotype of Δ*arc*A, Δ*fnr* and Δ*arc*AΔ*fnr*.** The overnight culture was pretreated with (A) sodium nitrate, (B) sodium nitrite, (C) sodium fumarate, (D) DMSO and then treated with 3% bile v/v for 6 h. O.D. was recorded after incubation at 37°C at 180 r.p.m. at 6 h. Data is representative of 3 individual experiments and presented as mean ± SD. *P* values were measured by two-way ANOVA using Turkey's multiple comparison test. $^{**}P < 0.01$, $^{****}P < 0.0001$.

**Supplementary Fig. 9**

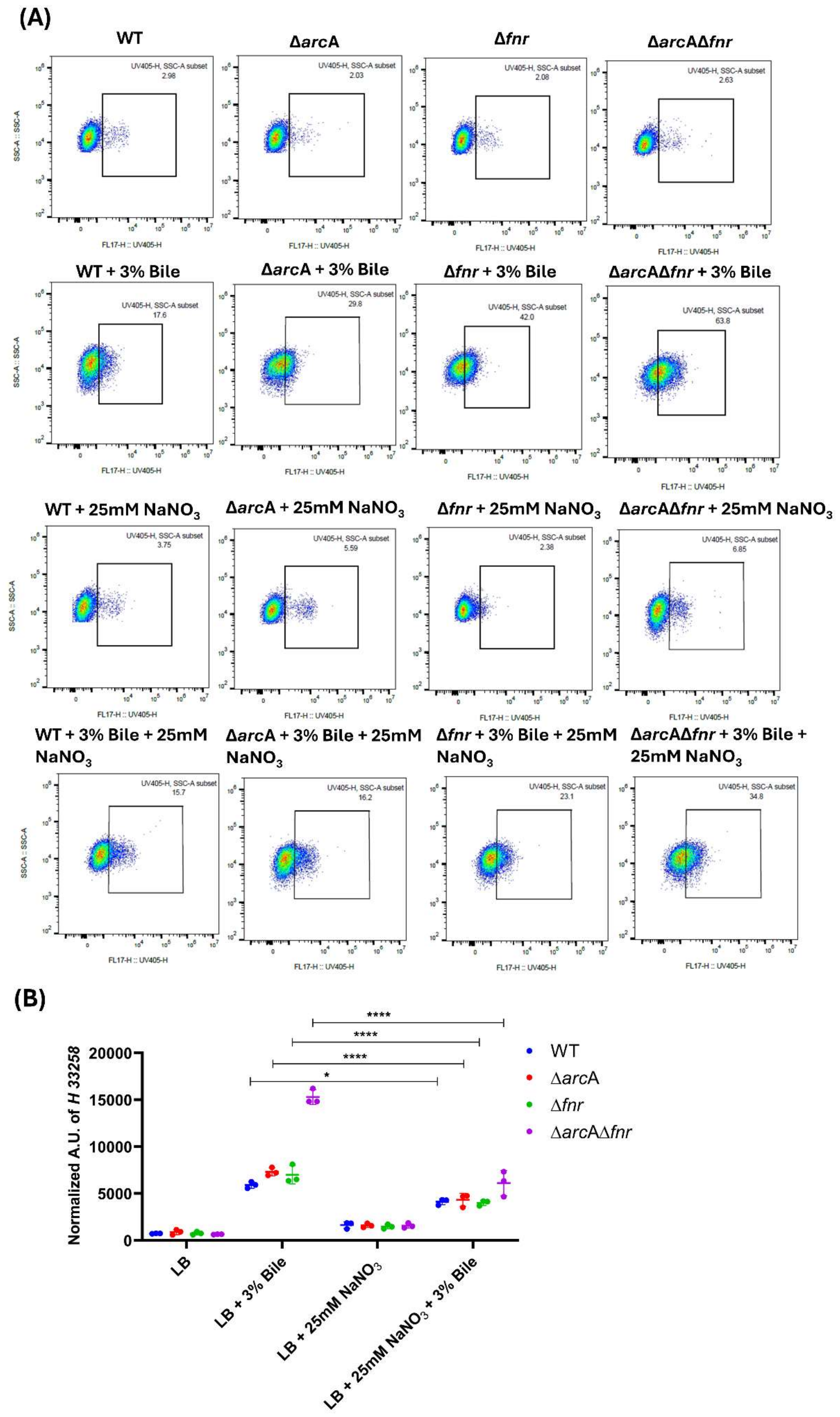

**Fig. S9 Pretreatment with sodium nitrate rescues the membrane damage in Δ*arc*A, Δ*fnr* and Δ*arc*AΔ*fnr*.** (A) Cells were pretreated with 25 mM $NaNO_3$ and stained with NPN post 6 h of bile treatment and analysed by flow cytometry. (B) Also, the pretreatment with sodium nitrate rescued the increased permeability in Δ*arc*A, Δ*fnr* and Δ*arc*AΔ*fnr*. Data is representative of 3 individual experiments and presented as mean ± SD. *P* values were measured by two-way ANOVA using Turkey's multiple comparison test. **P* < 0.1, ***P* < 0.01, ****P* < 0.001, *****P* < 0.0001.

**Supplementary Fig. 10**

**(A)**

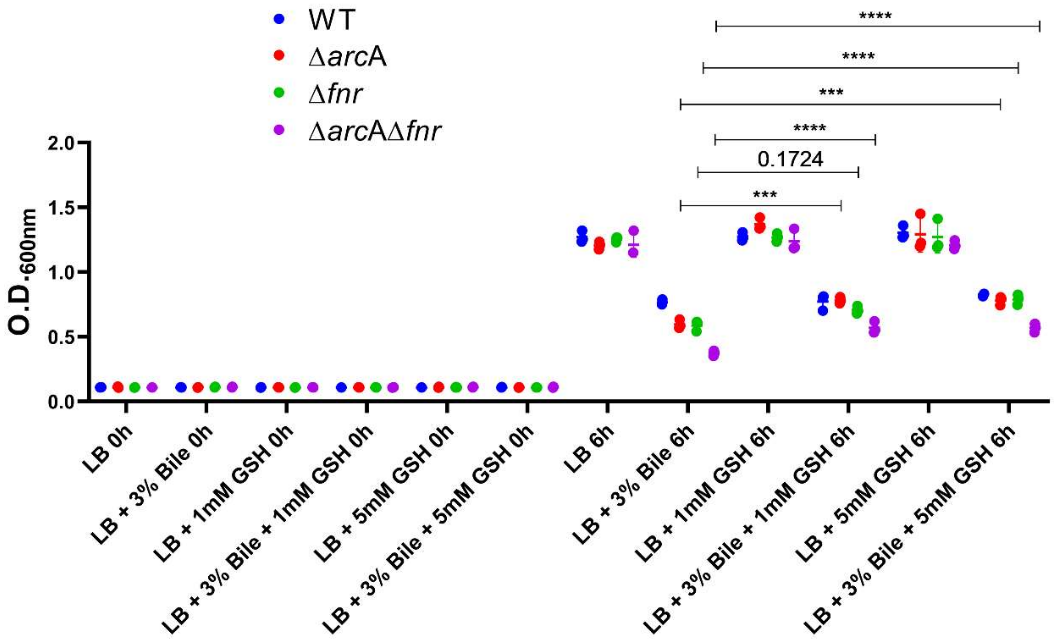


**(B)**

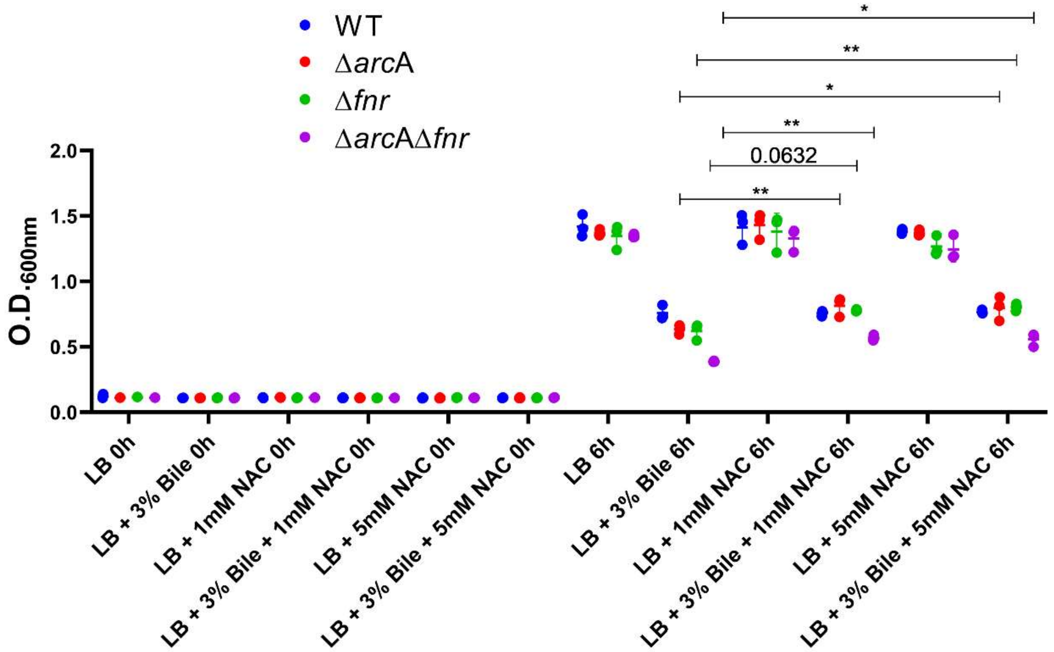


**Fig. S10 Pretreatment with GSH and NAC rescues the growth in Δ*arc*A, Δ*fnr* and Δ*arc*AΔ*fnr* upon bile exposure.** The overnight culture was pretreated with (A) GSH (B) NAC, and then treated with 3% bile v/v for 6 h. O.D. was recorded after incubation at 37°C at 180 r.p.m. at 6 h. Data is representative of 3 individual experiments and presented as mean ± SD. *P* values were measured by two-way ANOVA using Turkey's multiple comparison test. $^{**}P < 0.01$, $^{***}P < 0.001$, $^{****}P < 0.0001$.

### Supplementary Methodology

#### PCR confirmation of the single and double knockout strains

*Salmonella* Typhimurium 14028s was used as WT strain. A colony PCR was performed to identify WT, Δ*arc*A, Δ*fnr* and Δ*arc*AΔ*fnr* strains using the primers as mentioned in Table S3 to confirm *S*. Typhimurium. Negative control was used where no colonies were added.

#### Gene cloning

Using WT genomic DNA as a template and the primers indicated in the Table S4, PCR amplification of *arc*A and *fnr* was performed for cloning. The pQE60 plasmid was used to clone the genes *arc*A and *fnr* between the locations *Hin*DIII and *Bam*HI (New England Biolabs, Ipswich, MA, USA). The clones were confirmed by visualizing the insert release post restriction digestion and sanger sequencing (Aggrigenome, Kerala, India). The cloned constructs were transformed into Top10 competent cels. The positive clones: p*arc*A, p*fnr* and control vector pQE60 (VA) were transformed into *S*. Typhimurium WT, Δ*arc*A, Δ*fnr* and Δ*arc*AΔ*fnr* via electroporation, to generate the following strains; WT/VA, Δ*arc*A/VA, Δ*fnr*/VA, Δ*arc*AΔ*fnr*/VA, WT/p*arc*A, Δ*arc*A/p*arc*A, WT/p*fnr*, Δ*fnr*/p*fnr*, Δ*arc*AΔ*fnr*/p*arc*A, Δ*arc*AΔ*fnr*/p*fnr*.

#### RNA extraction

RNA extraction was performed as described previously (Singh et al., 2024). 50ml of overnight bacterial cultures of WT, Δ*arc*A, Δ*fnr* and Δ*arc*AΔ*fnr* were grown till $O.D._{600}$ of 0.3, followed by addition of 3% bile for 3 h. For the motility experiments the RNA was extracted post 6 h from the swimming agar plate from WT, Δ*arc*A, Δ*fnr* and Δ*arc*AΔ*fnr*. The methodology followed was as described before. The cells were washed twice with 1x PBS and resuspended in 1 ml TRIzol reagent (Ambion, Invitrogen, Carlsbad, CA, USA). They were allowed to lyse for 60 min at 1500 r.p.m. in shaking dry bath. 250 µl of chloroform (Sigma-Aldrich) was added to the cells and shaken vigorously for 1 min. The cells were centrifuged at 12000 g at 4°C for 15 min and the aqueous phase was collected in new tubes. The chloroform step was repeated as earlier followed by addition of 500 µl of 2-propanol (Merck, Steinheim, Germany) and kept in -20°C for 13 h. The RNA was pelleted at 15000 g at 4°C for 30 min and washed thrice with 75% ethanol. Pellet was air dried, and the RNA integrity was analysed on 1.5% agarose gel via electrophoresis.

**Gene expression analysis by qRT-PCR**

The RNA was processed with DNase I (New England Biolabs). cDNA was prepared as described previously (Singh et al., 2024). The table S5 lists the primers (Sigma Bangalore) used for qRT-PCR. 100 ng of cDNA samples, 250 nM of gene specific primers, and SYBR green master mix (BioRad, Hercules, CA, USA) were used to prepare PCR reactions, which were carried out in a Bio Rad CFX connect apparatus. The mean of three separate cDNA samples was used to determine each gene's expression level. Each cDNA sample and each gene were set up for experiment in triplicates. *gmk* and *gyr*B were used as reference genes and the normalized quantities of transcripts were estimated using the ΔΔCq method.

**Osmolar stress assay**

Overnight cultures of WT, Δ*arc*A, Δ*fnr* and Δ*arc*AΔ*fnr* strains were normalized to $O.D._{600}$ of 2. The normalized cultures of WT, Δ*arc*A, Δ*fnr* and Δ*arc*AΔ*fnr* strains were inoculated in 5ml LB and NaCl was added at 1 %, 2 %, 3 % and 4 % w/v concentration. The cultures were kept at 37°C for 6 h at 180 r.p.m. The $O.D._{600}$ was recorded using a Microplate reader (Tecan, Grodig, Austria) in a clear flat bottom 96 well culture plate (Tarsons, Kolkata, India).

**Temperature shift assay**

WT, Δ*arc*A, Δ*fnr* and Δ*arc*AΔ*fnr* strains were grown overnight and normalized to $O.D._{600}$ of 2. The cells were inoculated in 5ml LB for 6 h at 25° C, 37° C and 42° C. The $O.D._{600}$ was recorded using a Microplate reader (Tecan, Grodig, Austria) at 0 h, 3 h, 6 h, 9 h and 12 h in a clear flat bottom 96 well culture plate (Tarsons, Kolkata, India).

**Microaerobic growth analysis**

The overnight cultures of WT, Δ*arc*A, Δ*fnr* and Δ*arc*AΔ*fnr* were normalized to $O.D._{600}$ of 2. For the microaerobic trend the experiment was set up in a 15 ml falcon tube with LB media filled to brim followed by addition of bile to make up the final concentration of 3% v/v and was inoculated with the normalized cultures. The $O.D._{600}$ was recorded using a Microplate reader (Tecan, Grodig, Austria) in a clear flat bottom 96 well culture plate (Tarsons, Kolkata, India) for 0 h, 3 h, 6 h, 9 h and 12 h. C.F.U. was recorded by plating appropriate dilution of LB agar plate. For the microaerobic O.D. and C.F.U. the normalized increased volume of Δ*arc*A, Δ*fnr* and Δ*arc*AΔ*fnr* was added for comparison purpose and the $O.D._{600}$ was recorded. C.F.U. was recorded by plating appropriate dilution of LB agar plate. For the complemented strains the overnight grown cultures were normalized to $O.D._{600}$ of 2. The experiment was set up as described above followed by addition of bile to make up the final concentration of 3% v/v. The O.D.$_{600}$ was recorded using a Microplate reader (Tecan, Grodig, Austria) in a clear flat bottom 96 well culture plate (Tarsons, Kolkata, India) for 0 h, 3 h, 6 h, 9 h and 12 h. C.F.U. was recorded by plating appropriate dilution of LB agar plate.

**Colony morphology estimation**

Experiments were performed using the overnight grown cultures of WT/VA, Δ*arc*A/VA, Δ*fnr*/VA, Δ*arc*AΔ*fnr*/VA, WT/p*arc*A, Δ*arc*A/p*arc*A, WT/p*fnr*, Δ*fnr*/p*fnr*, Δ*arc*AΔ*fnr*/p*arc*A, Δ*arc*AΔ*fnr*/p*fnr* strains and normalized to O.D.$_{600}$ of 2. Appropriate dilutions were plated onto LB agar plates. These plates were allowed to be incubated for 36 h at 37°C unless otherwise mentioned (Chakraborty et al., 2025). Images for the colony morphology were obtained by a stereo microscope (*Nikon* SMZ745T MICAPS HPS3CMOS, Japan) at 20x.

**Hydrogen peroxide stress assay**

For complementation experiments, the overnight grown cultures of WT/VA, Δ*arc*A/VA, Δ*fnr*/VA, Δ*arc*AΔ*fnr*/VA, WT/p*arc*A, Δ*arc*A/p*arc*A, WT/p*fnr*, Δ*fnr*/p*fnr*, Δ*arc*AΔ*fnr*/p*arc*A, Δ*arc*AΔ*fnr*/p*fnr* strains were normalized to O.D.$_{600}$ of 2, 0.5mM $H_2O_2$ was added and incubated for 0 h and 6 h at 37°C. The *A* at 600 nm was recorded at 0 h and 6 h using a Microplate reader (Tecan, Grodig, Austria) in a clear flat bottom 96 well culture plate (Tarsons, Kolkata, India).

**Estimation of intracellular nitrite levels upon Glutathione and N-Acetyl Cysteine pretreatment.**

Overnight cultures of WT, Δ*arc*A, Δ*fnr* and Δ*arc*AΔ*fnr* was adjusted to O.D.$_{600}$ of 2. The cells were incubated with 1mM and 5 mM of Glutathione and N-Acetyl Cysteine and allowed to grow till O.D. 0.2. The cells were treated with 3% bile for 3 h. The cells were given three washes with 1 x PBS and proceeded towards sonication (QSonica Q700). Post cell lysis the cells were centrifuged at 13000 g for 15 min. 150 µl of supernatant was incubated with 100 µl of Griess reagent at room temperature in dark. O.D.$_{540}$ was recorded using a microplate reader (Tecan, Austria Gmbh).

**Estimation of intracellular ROS and bacterial growth upon Glutathione and N-Acetyl Cysteine pretreatment.**

Overnight cultures of WT, Δ*arc*A, Δ*fnr* and Δ*arc*AΔ*fnr* was adjusted to O.D.$_{600}$ of 2. The cells were incubated with 1mM and 5 mM of Glutathione and N-Acetyl Cysteine and allowed to grow till O.D. 0.2. The cells were treated with 3% bile for 6 h and *A* at 600 nm was recorded at 0 h and 6 h using a Microplate reader (Tecan, Grodig, Austria) in a clear flat bottom 96 well culture plate (Tarsons, Kolkata, India).

For ROS estimation the cells were incubated with 1mM and 5 mM of Glutathione and N-Acetyl Cysteine treated with 3% bile for 5 h. The cells were given three washes with 1 x PBS and incubated with 20 μM 2′,7′-Dichlorofluorescin Diacetate (DCFDA; Sigma-Aldrich) at 37 °C for 30 minutes. Post incubation with DCFDA dye the cells were washed twice with 1x PBS to remove excess unbound dye and 200 μl of the cell suspension was transferred to a 96 well plate. Fluorescence was measured using an Infinite 200 Pro plate reader (Tecan, Austria GmbH) at excitation/emission wavelengths of 485/535 nm. Fluorescence values were normalized to the $O.D._{600}$ of each sample.

**bisbenzimide H 33258 accumulation assay**

The overnight grown cultures were inoculated in 5 ml LB with and without treatment with 3% bile as described previously (Ray et al., 2019). The cells were allowed to grow for 5 h at 37 °C, 180 r.p.m. and then centrifuged at 5000 r.p.m. for 5 min. The cells were washed three times with 1x PBS and adjusted to $O.D._{600}$ of 0.1. 180 μl of culture was transferred to 96 well plate along with heat inactivated WT (90 °C, 10 min). The plates were then incubated with 2.5 μM bisbenzimide H 33258 (sigma) at 37 °C, in dark. Fluorescence values were recorded from the top of the well using excitation and emission wavelength of 355 and 460 nm, respectively. Appropriate control blanks and fluorescence values of unstained samples were subtracted from the samples. For imaging, the cells were treated with 3% bile for 5 h and normalized to 0.1 O.D. The normalized cells were treated with 2.5 μM bisbenzimide H 33258 (sigma) at 37 °C, in dark. Cells were washed three times with 1 x PBS and fixed with 4% PFA for 30 min in dark. The cells were washed three times with 1 x PBS to remove the excess dye and 70 μl of sample were drop casted on a clean glass slide and allowed to air dry for 30 min and sealed with cover slip. Images were acquired with the Zeiss LSM880 at 100x magnification (IISc, BC Central facility).

**Assessment of bile tolerance by pretreatment with various electron acceptors**

Overnight cultures of WT, Δ*arc*A, Δ*fnr* and Δ*arc*AΔ*fnr* was adjusted to $O.D._{600}$ of 2. The cells were inoculated in 5 ml LB with and without supplementation with sodium nitrite, sodium fumarate and DMSO in a dose dependent manner. The cells were allowed to grow till $O.D._{600}$

of 0.2 and then treated with bile for 6 h at 37 °C, 180 r.p.m. O.D.$_{600}$ was recorded using a microplate reader and appropriate dilutions were plated in LB agar plate to calculate the CFUs.
